# Genome-wide comprehensive analysis of miRNAs and their target genes expressed in resistant and susceptible *Capsicum annuum* landrace during *Phytophthora capsici* infection

**DOI:** 10.1101/2021.04.13.439754

**Authors:** Tilahun Rabuma, Om Prakash Gupta, Vinod Chhokar

**Affiliations:** Department of Bio and Nano Technology, Guru Jambheshwar University of Science and Technology, Hisar, Haryana, India; Division of Quality and Basic Sciences, ICAR-Indian Institute of Wheat and Barley Research, Karnal-132001, Haryana, INDIA

**Keywords:** C. annuum, P. capsici, miRNA, defence response, transcriptome, NGS

## Abstract

MiRNAs regulate plants responses to fungal infection and immunity by modulating the gene expression. Despite extensive works on miRNA’s role during plant-fungus interaction, work in *Capsicum annuum-Phytophthora capsici* pathosystem is limited. Therefore, in the current study, genome-wide known and novel miRNAs were identified in two contrasting chilli pepper landraces, *i.e.* GojamMecha_9086 (resistant) and Dabat_80045 (susceptible) during *P. capsici* infection. The small RNA deep sequencing resulted in 79 known miRNAs corresponding to 24 miRNAs families and 477 novel miRNAs along with 22,895 potential targets, including 30 defence-related genes against *P. capsici* infection. The expression analysis of ∼29 known & 157 novel miRNAs in resistant and 30 known and 176 novel miRNAs in susceptible landrace revealed differential accumulation pattern. RT-qPCR of a set of 8 defence related miRNAs representing 4 novel (Pz-novel-miR428-1, Pz-novel-miR160-1, Pz-novel-miR1028-1, Pz-novel-miR204-1) and 4 known (Pz-known-miR803-1, Pz-known-miR2059-1, Pz-known-miR2560-1, Pz-known-miR1872-1) revealed differential accumulation pattern in both resistant and susceptible landrace. Additionally, validation of 8 target genes of corresponding miRNAs using RA-PCR, which as good as 5’ RLM-RACE, revealed an inverse relation with their corresponding miRNAs suggesting their key role during disease response. This study provides comprehensive genome-wide information about the repertoire of miRNAs and their target genes expressed in resistant and susceptible chilli pepper landrace, which can serve as a valuable resource for better understanding the post-transcriptional regulatory mechanism during *C. annuum* - *P. capsici* pathosystem.

## Introduction

Plants exposed to stress use multiple gene regulatory mechanisms, including post-transcriptional regulation of gene expression, to restore and re-establish cellular homeostasis. During the last two decades, microRNAs (miRNAs) have emerged as one of the ubiquitous and critical regulatory molecules during the entire life cycle of nearly all eukaryotes (Bartel, 2004). The use of miRNAs for trait improvement has been successfully demonstrated wherein overexpression of *Os*miR397 enhanced rice yield by increasing grain size and promoting panicle branching (Zhao *et al*., 2013). The latest release of miRBase(v22) reported to contains 38,589 hairpin precursors and 48,860 mature microRNAs sequences from 271 organisms showing a continuous increase in the miRNA pool (Griffiths *et al*., 2019). With the advancement in high throughput technology and rapidly growing bioinformatics, several workers have identified miRNAs, which play a critical role during disease pathogenesis upon fungal invasion in plants by interacting with several classes of resistance genes (Yin *et al*., 2012; Chen *et al*., 2012). The data generated by the high throughput sequencing platforms have uncovered the potential role of miRNAs in plants during fungal invasion (Gupta *et al*., 2014). It has been observed that the host miRNAs, along with their targets either by up or down-regulation, participate in disease modulation in plants upon fungal infection (Zhao *et al*., 2012; Gupta *et al*., 2012). For instance, miR1138 was found to be highly accumulated in wheat infected with *P. graminis f.sp. tritici* (62G29-1) (Gupta *et al*.,2012). Din *et al*. (2016) had profiled and characterized conserved miRNAs and their targets in chilli (*Capsicum annuum* L.) using ESTs. Yang and co-workers (Yang *et al.,* 2021) investigated the heterosis among three cross combinations of *C. annuum* varieties by integrating mRNA and miRNA profiling. Zhang *et al*. (2020) identified miRNAs and their targets involved during anther development in CMS-line N816S and its maintainer line Ning5m pepper (*C. annuum*). Shin and co-workers (Shin *et al*., 2013) computationally predicted miRNA target in hot pepper (*C. annuum*). The miRNA targets of pepper and their regulatory networks were explored through techniques of MiRTrans (Trans-omics data) (Zhang *et al*., 2017). Liu and co-workers (Liu *et al.,* 2017) identified and validated miRNAs involved in fruit development and fruit ripening in hot pepper. However, despite these reports, there is no work on identifying and characterizing miRNAs and their targets during *C. annuum L* – *P. capsici* pathosystem. Therefore, understanding miRNA expression and regulation during *C. annuum* - *P. capsici* pathosystem will help in a better understanding of the molecular and biochemical pathways associated with defence response in chilli pepper.

Pepper (*Capsicum annuum* L.) is one of the most economically important vegetable crops in the world (Catalan *et al*., 2020) owing to its diverse usages, including spice, either fresh or in powder form (El-Ghoraba *et al*., 2013), drug, condiment, ointment (Chamikara *et al*., 2016). On the other hand, its productivity is highly affected by both abiotic and biotic factors. Among biotic factors, the fungal pathogens, especially *P. capsici* imposes severe threats to its production as it can cause losses up to 100% (Babadoost, 2004). The *P. capsici* is a soil-born pathogen triggering diseases like root and crown rot, leaf late blight stems blight, and fruits (Mao *et al*., 1998). Understanding the molecular mechanism and developing resistant genotypes is a sustainable approach to tackle it as control by chemical and cultural practices remains a challenge (Grumet *et al*., 2009). Identification and characterization of novel and known miRNAs in chilli pepper in response to *P. capsici* shall widen our current understanding of these miRNAs’ role in modulating the molecular and biochemical pathways with disease resistance.

Four small RNA libraries were prepared from control and infected leaves of *P. capsici* resistant and susceptible chilli pepper landraces to fill this gap. The two contrasting chilli pepper landraces, *i.e.* GojamMecha_9086 (resistant) and Dabat_80045 (susceptible), were selected based on our earlier report (Rabuma *et al.,* 2020). The libraries represented small RNAs expressed during control and infection of both resistant and susceptible landraces. We utilized genome-wide sequencing of small RNAs to identify known and novel expressed miRNAs in resistant and susceptible landraces. Target genes of these miRNAs were also detected from our previous transcriptome data (under review) of the same samples. Besides, expression analysis of 8 key miRNAs, along with cleavage/inhibition pattern of their eight corresponding target genes associated with defence response against fungal invasion and immunity, were also validated using RT-qPCR and RA-PCR, respectively.

## Materials and Methods

### Plant material, fungal strain, and sample preparation

Based on disease score data published earlier (Rabuma *et al*., 2020), two contrasting landraces of chilli pepper *(C. annuum), i.e.* GojamMecha_9086 (resistant) and Dabat_80045 (susceptible), were selected for small RNA sequencing and identification of miRNA to unravel the defence mechanism operating during *C. annuum – P. capsici* pathosystem. The details of germination, growth conditions, fungal strain, sporulation and inoculation are the same as described earlier (Andrés Ares *et al*., 2005; Rabuma *et al*., 2020). After five days post-inoculation (dpi), samples were harvested from three biological replications of control and infected leaves (4^th^ and 5^th^ leaves) of GojamMecha_9086 and Dabat_80045 representing resistant control (RC), resistant infected (RI), susceptible control (SC) and susceptible infected (SI). The leaf samples were washed and frozen in liquid nitrogen and kept at −80 °C in a deep freeze refrigerator. The total RNA was isolated using ZR Plant RNA Miniprep™ Kit by Zymo Research as per manufacturer instruction. Total RNA was isolated separately from control and infected leaf tissues. The quality and quantity of isolated RNA were checked on 1% denaturing agarose gel and Nanodrop spectrophotometry (Fluorometer, DS-11 FXT, Denovix), respectively. Small RNA enriched samples were sequenced using the Illumina NextSeq500 sequencing platform.

### Small RNA library preparation and Illumina sequencing

The small RNA sequencing libraries were prepared from quality checked (QC) passed total RNA using Illumina TruSeq small RNA library preparation kit (Cat. No-RS-200-0024) according to the manufacturer instruction. High-quality small RNA reads were obtained from raw reads by filtering out poor quality reads and removing adapters contamination using Cutadapt (v 1.16) by considering parameters, *i.e*. any reads with < 10% quality threshold (QV) < 20 Phred score were removed, reads with adaptors were discarded, a quality with a minimum overlap between the adapter and read was 5bp, reads with ambiguous base pair(N) and reads length below 15bp were trimmed off. After removing the adapter sequences, the quality filtering analysis was performed to trim the low-quality terminal bases using the Trimmomatic (v. 0.38) in a single end mode with a minimum read length of 15bp and base quality cut-off of 20. The quality box plot graph was generated to visualize the quality score of each base with respect to its base position. Quality filtering was performed using the fastq quality filter program, and a quality cut-off (q) of 20 and a minimum percentage (P) value of 100 was assigned for pre-processing. A quality-filtered reads were formatted into a non-redundant FASTA format for the unification of reads. Occurrences of each unique sequence reads were counted as sequence tags. The number of reads for each unique tag reflects the relative expression level of that miRNA. The bioinformatics workflow for Small RNA sequencing and data analysis is shown in supplementary Fig. S1. The sequence data have been submitted to NCBI. The BioProject ID is….

### Identification of known and novel miRNAs

The clean read tags from each library were blasted against the customized Rfam database (http://www.sanger.ac.uk/software/Rfam), and any small RNA read tag mapped to the Rfam database were filtered out and excluded from further analysis by considering the parameters, *i.e.* BlastN variant (blastn-short), word size (7bp), low complexity filtering (off), identity (100%), query coverage (100%), mismatch and gaps (not allowed). The reads tags that were not mapped to Rfam were further mapped to known repeat sequence database (Repbase (v 22)) using NCBI blast and were excluded from further analysis if not satisfying the stringent criteria as mentioned above. Read length filtration analysis was performed and reads with length >34nt excluded while <34 nt. (nucleotides) were considered clean putative miRNA reads. These putative miRNA reads were subsequently used for the identification of known and novel miRNA. The putative miRNA reads were mapped to the *Capsicum annuum* (Pepper Zunla v1.0) Genome (GCA_000710875.1) obtained from the NCBI Genome database. Precursor sequences were identified using the tool miRCat from UEA small RNA workbench (v3.2). The precursor stability was analyzed by estimating the MFE (minimum free energy), and only miRNAs observed on stable precursor sequences were reported (Tiwari *et al*., 2020). To identify known miRNAs and assign miR family, all the unique sequences were BLASTED against the known plant miRNA database (http://www.mirbase.org, release v21). Based on the assigned miRbase ID, it was segregated as the known and novel miRNAs.

### Differential analysis and target prediction

The sample wise tag counts were used for expression profiling of known and novel miRNA, and miRNAs with at least five reads in any of the four samples were considered for the differential analysis. TPM normalization algorithm was done, and the TPM values were then logged (base 2) transformed to get fold expression analysis. Finally, fold-change values were calculated by subtracting respective fold expression values. Fold change value of above zero was considered as up-regulated, and below zero was considered as down-regulated. For predicting a potential target, conserved and non-conserved miRNAs of *C. annuum* were used query sequences for blast searches against our *C. annuum* transcriptome sequences (deposited in NCBI database under accession number RC: SAMN16251280; RI: SAMN16251798; SC: SAMN16251797; SI: SAMN16251799 on SRA at the link https://www.ncbi.nlm.nih.gov/biosample/16251797 using the methods in Wan *et al*., (2012).

### Validation of microRNAs sequencing by qRT-PCR

We utilized the same RNA used for small RNA-seq for qPCR-based validation of selected miRNAs along with their corresponding target genes. The cDNA was synthesized using Mir-X^TM^ miRNA First-Strand synthesis kits according to the manufacturer’s protocol. For validation of differentially expressed genes, eight miRNAs representing four novel and four known miRNAs and their eight corresponding target genes associated with modulating the disease pathogenesis during *P. capsici* infection were selected. The list of miRNA specific forward primers is given in Supplementary Table S1. RTQ-UNIr was utilized as a universal reverse primer. To further measure and validate the corresponding target genes’ expression levels, regional amplification quantitative RT-PCR (RA-PCR) assay was performed. The RA-PCR was developed to monitor the miRNA-directed cleavage of mRNAs (Oh *et al*., 2008). MicroRNA mediated cleavage of the mRNA transcripts’ target site leads to a decrease in RT-PCR accumulation of any fragment present upstream of the target site (Navarro *et al*., 2006). The reverse transcription of the miRNA-cleaved mRNA will not generate a cDNA beyond the cleaved site. Therefore, the cDNA segment present upstream of the cleaved site can be expected to be less abundant than a segment present downstream of the cleaved site. RA-PCR has added benefits of including control reaction in the same experiment while other technique such as 5′ rapid amplification of cDNA ends (RACE) 5′ RACE requires separate reaction and could be skewed by the presence of false positives. For each of the eight target genes, three sets of primer (RA-5’ region, RA-middle region and RA-3’ region) were designed using Express 3.0.1 primer software (Fig 5; Supplementary Table S2). Actin-7 (XM_016695142.1) was used as an endogenous control gene for data normalization (Gupta *et al*., 2017).

RT-qPCR reaction was performed in three technical replication from each of the three biological replication using SYBR Green JumpStart Taq ReadyMix 2x on Applied Biosystems™ StepOne™ Real-Time PCR System. RT-qPCR reaction was set in a 20µl reaction volume containing 10µl SYBR Green JumpStart Taq ReadyMix (2x), 2µl forward primer (10 μM), 2μl RTQ-UNI reverse (10 μM), 4μl cDNA and 2μl of nuclease-free water of molecular biology grade with the programme of 94 °C and denatured for 2 min, then primary reaction (denaturation at 94°C for 15 seconds, annealing at 55-60°C for 60 seconds, and template extension at 72°C for 15 seconds) for 40 cycles. The *actin-7-like* gene (Sequence ID: XM_016695142.1) and U6 snRNA (snRNA) (GenBank: Z17301.1) was used to normalize target and miRNA gene expression, respectively. The relative expression level was calculated using the 2^-ΔΔCt^ method (Livak and Schmittgen, 2001). The standard error in expression levels of the three biological replications in the figures was denoted by error bars, and the results were expressed as mean value ± SD (Hao *et al*., 2016).

## Results

### Sequencing of small RNA

To identify chilli pepper miRNAs induced by *P. capsici* infection, we analyzed small RNA sequencing data from control and infected chilli pepper leaf tissues of resistant (GojamMecha_9086) and susceptible (Dabat_80045) chilli pepper landrace. Illumina sequencing generated a total of ∼70 million high-quality clean reads representing 19, ∼7, ∼22 and ∼20 million from RC, RI, SC, and SI libraries, respectively, after removing low-quality sequences and adapters (Table 1). After filtering the low-quality tags (removing non-coding small RNAs), trimming 3’ adaptors and removing contaminants, ∼8 million clean reads of 18–24 nucleotide lengths were obtained for downstream analysis of the miRNAs (Table 1). The sequencing data’s reliability was ensured by average quality (Q30 score) of 93.45%, ranging from 93.11% to 93.87% across the libraries (Table 1).The miRNAs’ length distribution showed the highest abundance of reads of 24 nucleotides followed by 21 nt., 22ntd and 23 nt. in all the four libraries with the maximum in RC and SC compared to RI and SI (Fig. S2).

**Table 1.**
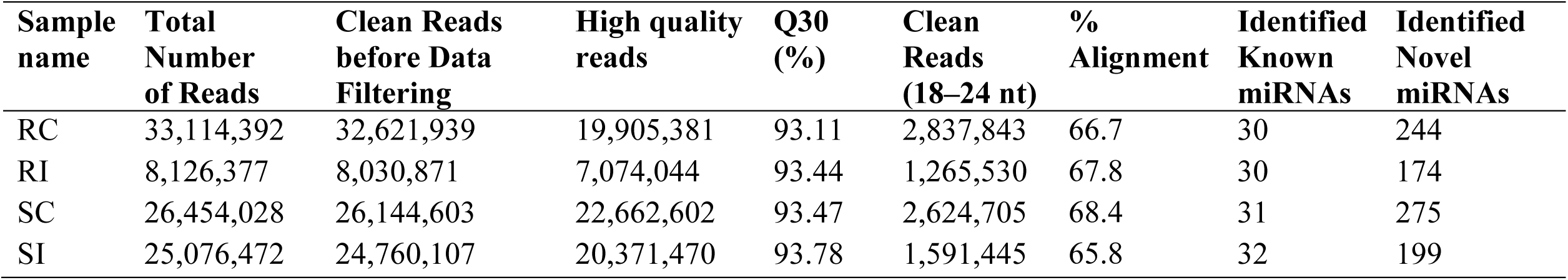
Summary of small RNA sequencing and analysis in control and *P. capsici* infected libraries of *C. annuum* L.

| **Sample name** | **Total Number of Reads** | **Clean Reads before Data Filtering** | **High quality reads** | **Q30 (%)** | **Clean Reads (18–24 nt)** | **% Alignment** | **Identified Known miRNAs** | **Identified Novel miRNAs** |
| --- | --- | --- | --- | --- | --- | --- | --- | --- |
| RC | 33,114,392 | 32,621,939 | 19,905,381 | 93.11 | 2,837,843 | 66.7 | 30 | 244 |
| RI | 8,126,377 | 8,030,871 | 7,074,044 | 93.44 | 1,265,530 | 67.8 | 30 | 174 |
| SC | 26,454,028 | 26,144,603 | 22,662,602 | 93.47 | 2,624,705 | 68.4 | 31 | 275 |
| SI | 25,076,472 | 24,760,107 | 20,371,470 | 93.78 | 1,591,445 | 65.8 | 32 | 199 |

### Identification and expression analysis of known miRNAs during *C. annuum – P. capsici* interaction

The clean reads of small RNAs (18–24 nt.) obtained after filtration were mapped against *Capsicum annuum* (Pepper Zunla v1.0) genome (GCA_000710875.1), and known miRNAs were searched against the miRNA database (miRBase 21 released) (Zhang *et al*., 2020). The small RNA sequence analysis obtained 79 known miRNAs corresponding to 24 miRNAs families in the four-leaf sample libraries of chilli pepper (Table 1). The highest number of known miRNAs (32) was identified in the infected leaf tissue of susceptible landrace Dabat_80045, SI, whereas the lowest number (30) was identified in resistance control (RC) and resistance infected (RI) leaf (Table 1). Amongst all, thirty commonly expressed known miRNAs were identified in RI and SI leaf, whereas 29 known miRNAs were commonly detected in RC and SC samples (Fig. 4 A; Supplementary Table S3). The differential miRNA expression analysis of know miRNA revealed down-regulation of 4 and 5 miRNAs in RC vs RI and SC vs SI, respectively, whereas up-regulation of 25 miRNAs in RC vs RI and SC vs SI, respectively (Table 3). The most abundant known miRNA families in all the four libraries were Pz-known-miR356-1(mir482), Pz-known-miR1099-1(mir159), Pz-known-miR745-1(mir166), Pz-known-miR2908-1(mir396), and Pz-known-miR3049-1(mir482). Pz-known-miR3686-1(mi395) was abundantly detected only in SI, while Pz-known-miR1186-1(mir156) in only SC leaf. Among all the known miRNA families, Pz-known-miR2908-1(mir396) was most abundant in all the four samples with the highest accumulation in resistant control landrace GojamMecha_9086. Expression analysis of miRNA families further revealed the highest number of miRNA families mir159 (Pz-known-miR1099-1) in RC (36427) sample while lowest in RI (11157) sample. Compared to control conditions, different miRNA families’ expression behaviour was greater during *P. capsici* infection. Additionally, the secondary structure of the four most abundant known miRNA families and novel miRNAs were given in Fig. 1A-D and 2A-D, respectively.

**Fig. 1.**
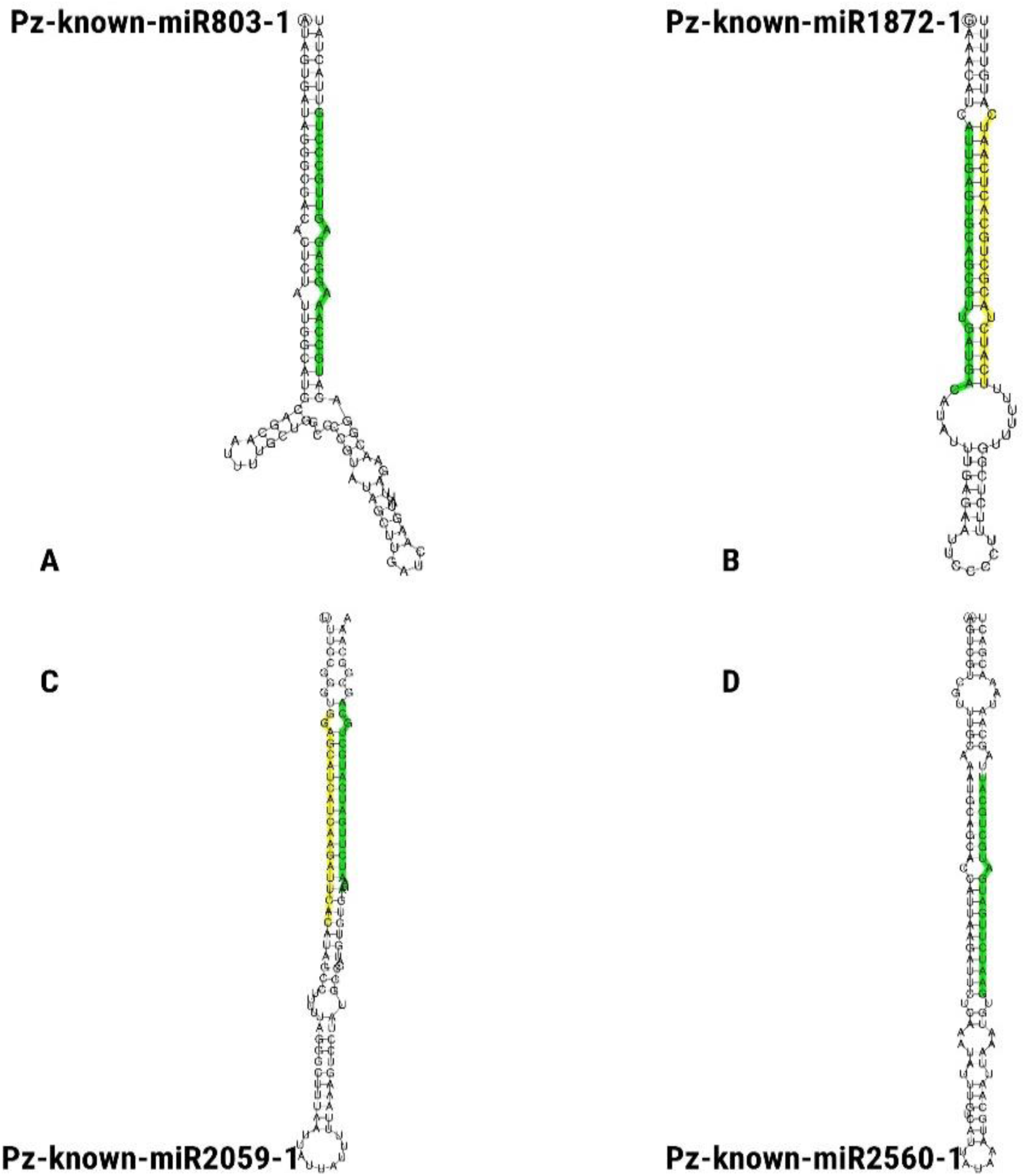
Secondary structure of known miRNA identified from small RNA sequencing of *C. annuum* during infection with *P. capsici*: A) Pz-known-miR803-1 (can-mir399), B) Pz-known-miR1872-1 (can-mir397), C) Pz-known-miR2059-1 (can-mir172), D) Pz-known-miR2560-1 (can-mir172) used in the validation of RT-qPCR (the green colour shows the matured miRNA sequence site).

**Fig. 2.**
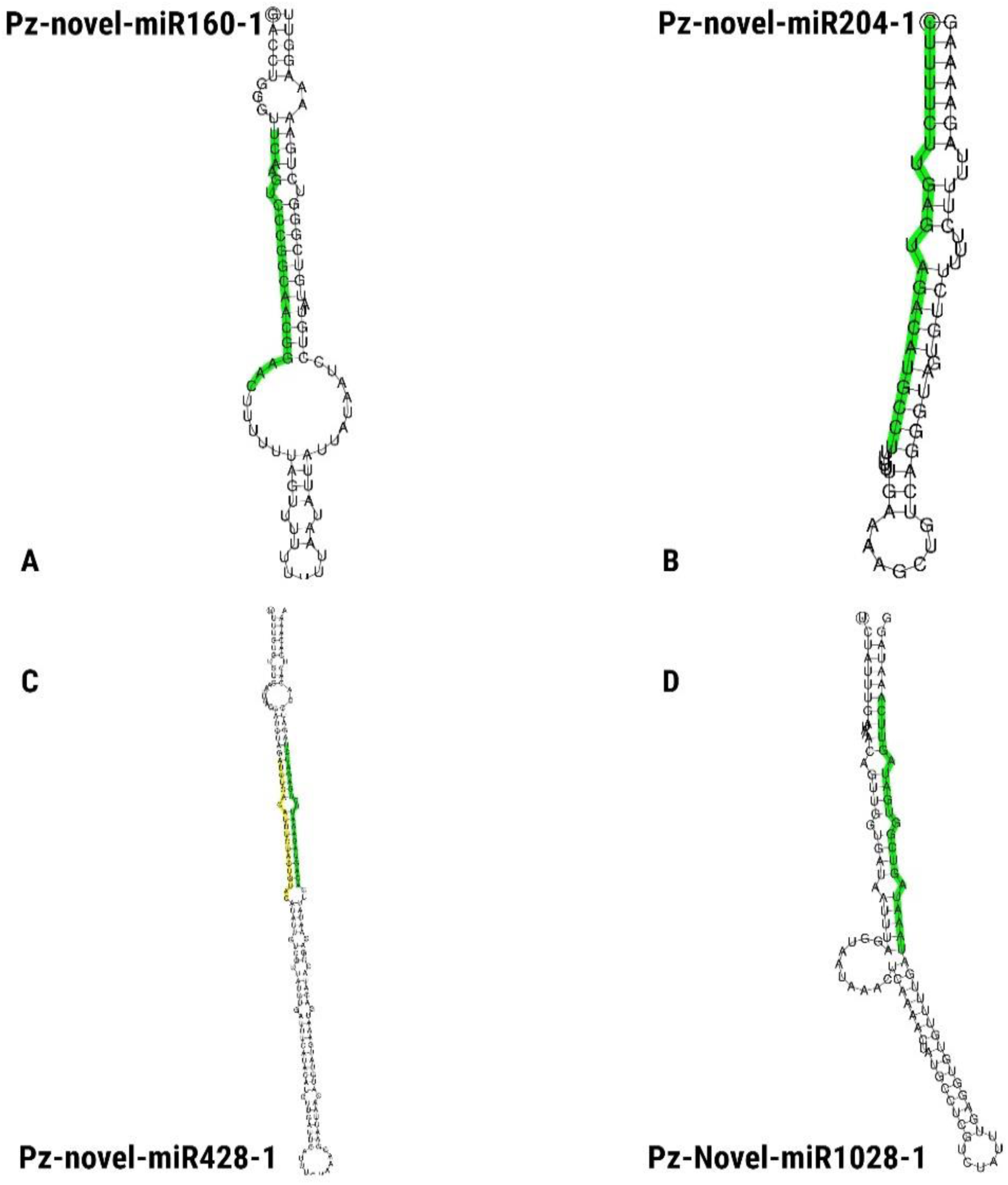
Secondary structure of novel miRNAs identified from transcriptome sequence of *C. annuum* during infection of *P. capsici* and the matured miRNAs sequence marked with green colour: A) Pz-novel-miR160-1, B) Pz-novel-miR204-1, C) Pz-novel-miR428-1, and D) Pz-novel-miR1028-1

### Identification, distribution and expression analysis of novel miRNAs during *C. annuum – P. capsici* interaction

The miRNA sequencing of RC, RI, SC, SI leaf samples predicted a total of 477 novel miRNA with the highest in SC (∼275), followed by RC (∼244), SI (∼199) and RI (∼174) (Table 1). The differential expression analysis of novel miRNA revealed down-regulation of 6 and 32 miRNAs in RC vs RI and SC vs SI, respectively, whereas up-regulation of 151 and 144 miRNAs in RC vs RI and SC vs SI, respectively (Table 3). The most abundant expressed novel miRNA in all the four libraries were Pz-novel-miR3731-1, Pz-novel-miR1981-1, Pz-novel-miR3509-1, Pz-novel-miR2406-1, Pz-novel-miR227-1, and Pz-novel-miR1344-1 (Supplementary Table S4). The Pz-novel-miR3731-1 was the highest abundant in number all four sample with (177934) in SC leaf sample while the lowest in RI (19510) sample (Supplementary Table S4). Most of the known and novel miRNA was found to be common in all the samples, while some of miRNAs were specifically expressed to specific leaf sample. The length of the consensus matured miRNA sequence of the novel miRNAs was in the range of 19 to 24 nt., of which the majority were 24 nt. (Fig. S2; Table 2). The number of predicted novel miRNAs were more in the control leaf than the infected leaf in the two genotypes (Table 1). Chromosomal distribution analysis of the novel miRNAs showed the involvement of all the chromosomes with the maximum on chromosomes 1 (50) and lowest on chromosome 4 (31), 8(30) and 12 (30) (Fig 3). Additionally, the secondary structure of the novel miRNAs was also predicted, and a representative of the four most abundant novel miRNAs showing mature miRNAs located on either arm of the secondary hairpin structures are shown in Fig. 2A-D). Also, we observed quite a good negative free energy of hairpin structure ranging from −131.29 to −15.3 kcal/mol (Table 2), which is comparable with previous reports in hot pepper (−104.85 kcal/mol), Arabidopsis *thaliana* (−59.5 kcal/mole) and *Oryza sativa* (−71.0 kcal/mol) (Shin *et al*., 2013). A total of 29 known and novel miRNAs (Supplementary Table S5) were identified as potential players regulating target genes associated with defence response against *P. capsici* infection. Out of that, 18 of these miRNAs were identified as novel miRNAs associated with defence response during *P. capsici* infection.

**Fig. 3.**
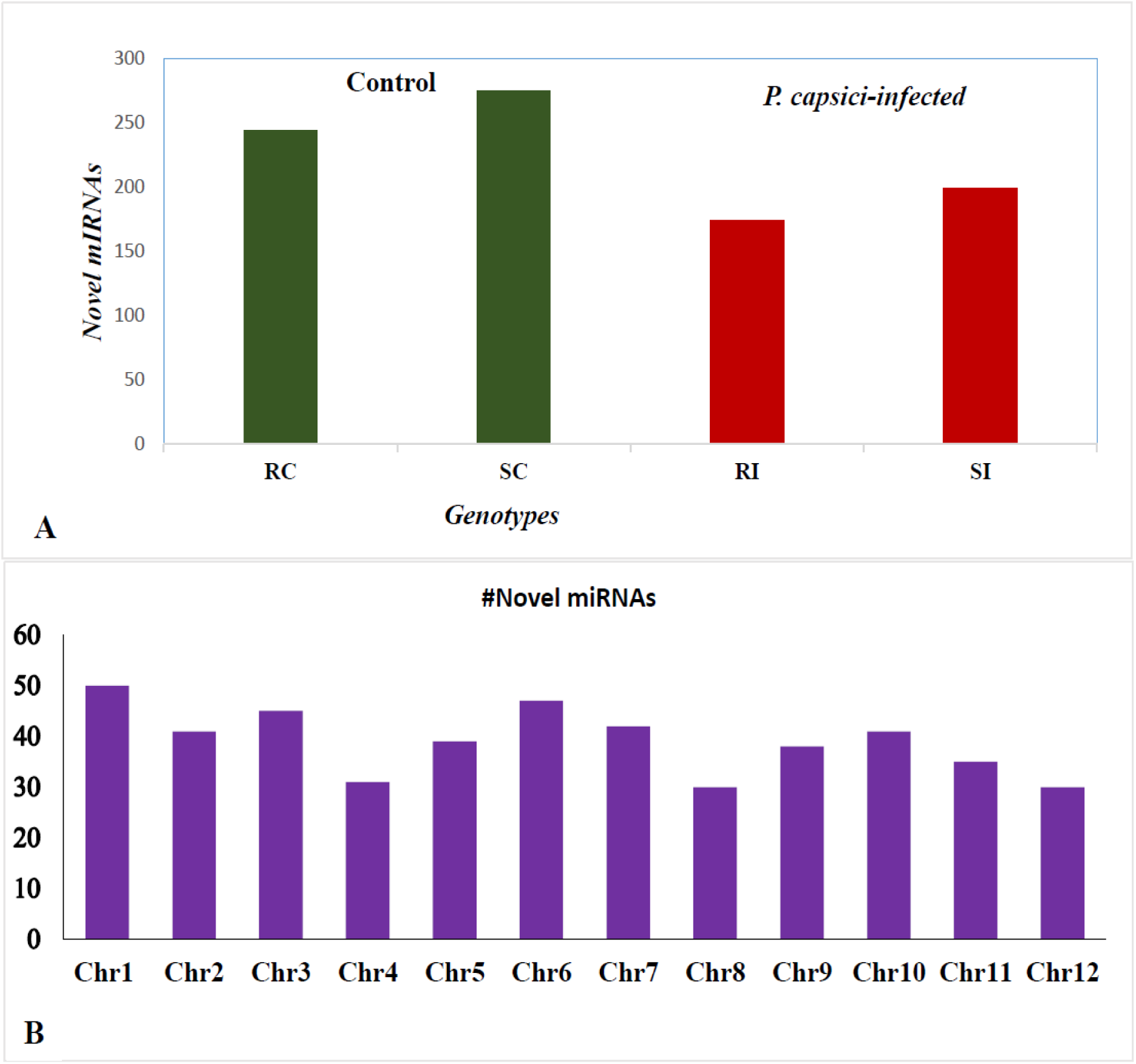
Novel miRNA identified in *C. annuum* A) Distribution of novel miRNAs in control (dark green colour bars) and in infected (red colour bars) samples of *C. annuum*; B) Distribution of novel miRNAs in different chromosomes of the *C. annuum* genome.

**Table 2.**
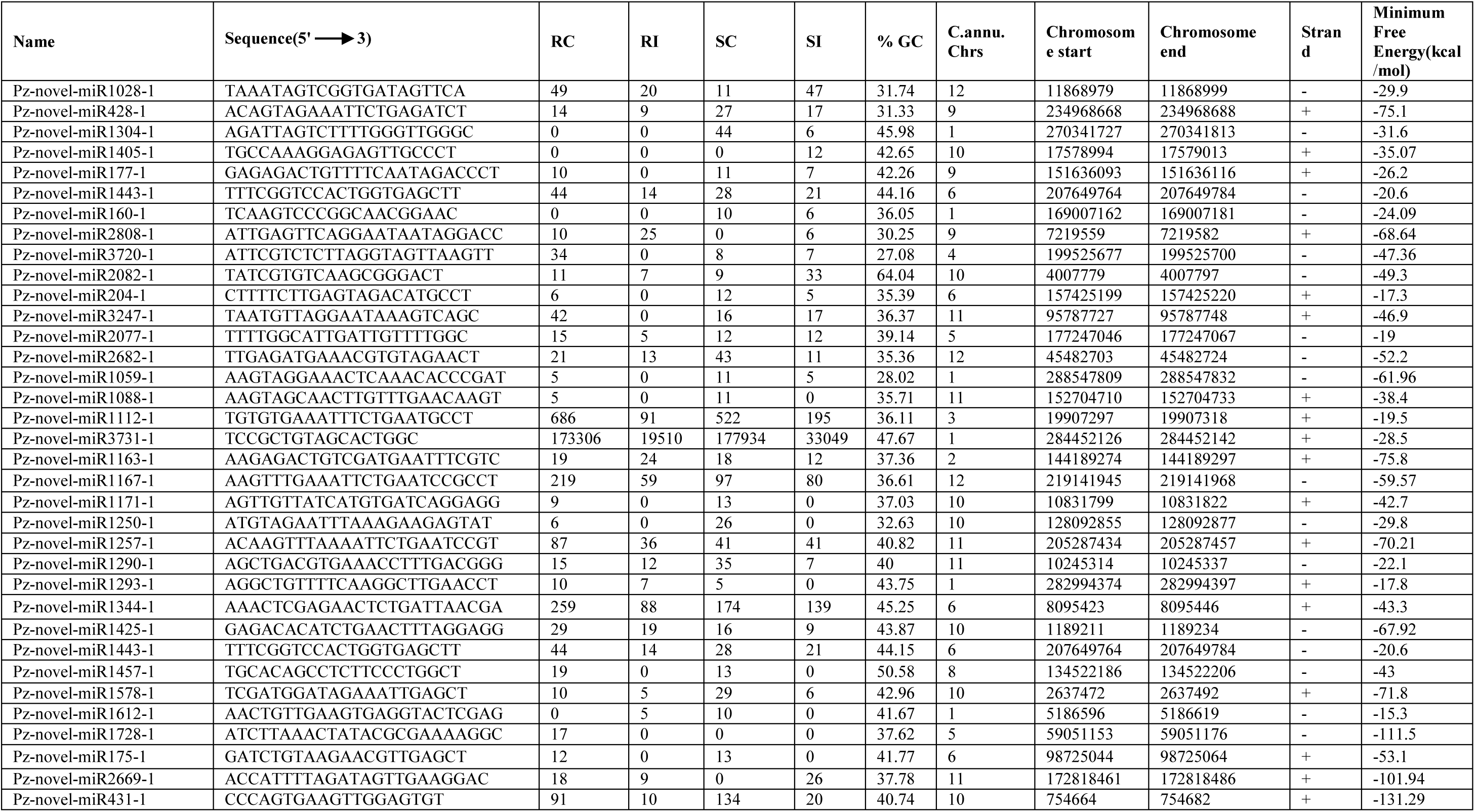
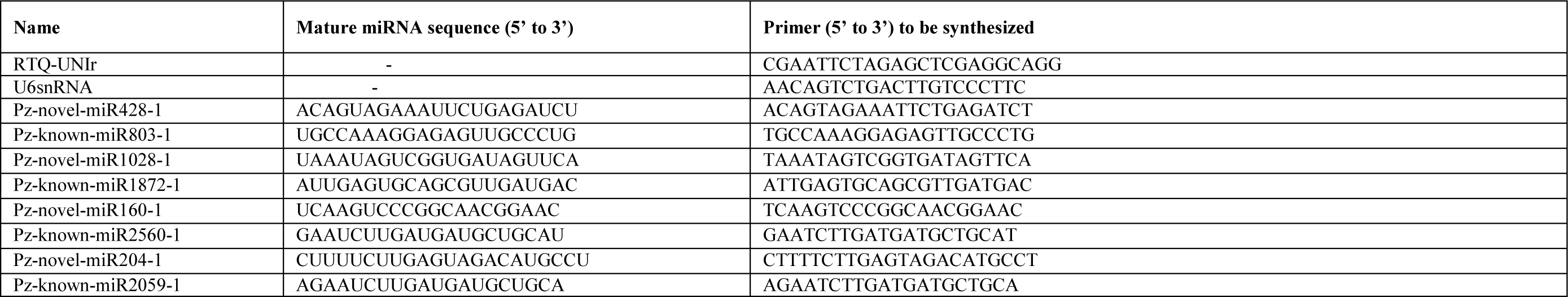
Some identified novel miRNAs and their abundance with minimum free energy across the four-leaf library of *C. annuum* under *P. capsici* infection.

### Comparative expression analysis of miRNAs in control and infected chilli pepper samples

The expression of miRNAs in control and infected leaf were compared for each of the two contrasting chilli pepper landraces. The expression of favoured miRNAs under the control and infected conditions were categorized along with the commonly expressed miRNAs (Fig. 4 and Supplementary Table S3). The result indicated fewer novel and known miRNAs in infected leaf tissue than control (Fig. 4). Further, these miRNAs were analyzed to identify the common miRNAs expressed under infected and control tissue of both the landraces. The result indicated that 87, 17, 98, and 22 novel miRNAs were explicitly expressed in RC, RI, SC, and SI, respectively (Fig.4). Among them, Pz-novel-miR2320-1 (∼7.9 fold) and Pz-novel-miR2429-1 (∼7.8 fold) were highly explicitly expressed in infected RI and SI library, respectively. Additionally, 177 novel miRNAs were commonly expressed between SC and SI, whereas 157 novel miRNAs were shown commonly expressed between RC and RI libraries. Moreover, 29 known miRNAs were commonly expressed between RC and RI libraries, while 30 known miRNAs were commonly expressed between SC and SI libraries. However, Pz-known-miR2226-1(mir398) (∼5.5 fold) and Pz-known-miR504-1(mir1446) (∼7.0 fold) were explicitly expressed in RC and RI leaf, respectively, while Pz-known-miR1186-1 (mir156) (∼5.2 fold) and two known miRNAs, i.e. (Pz-known-miR2226-1(mir398) (∼8.0 fold) and Pz-known-miR3686-1(mir395) (∼7.4 fold)) were expressed explicitly in SC and SI leaf, respectively (Fig 4).

**Fig. 4.**
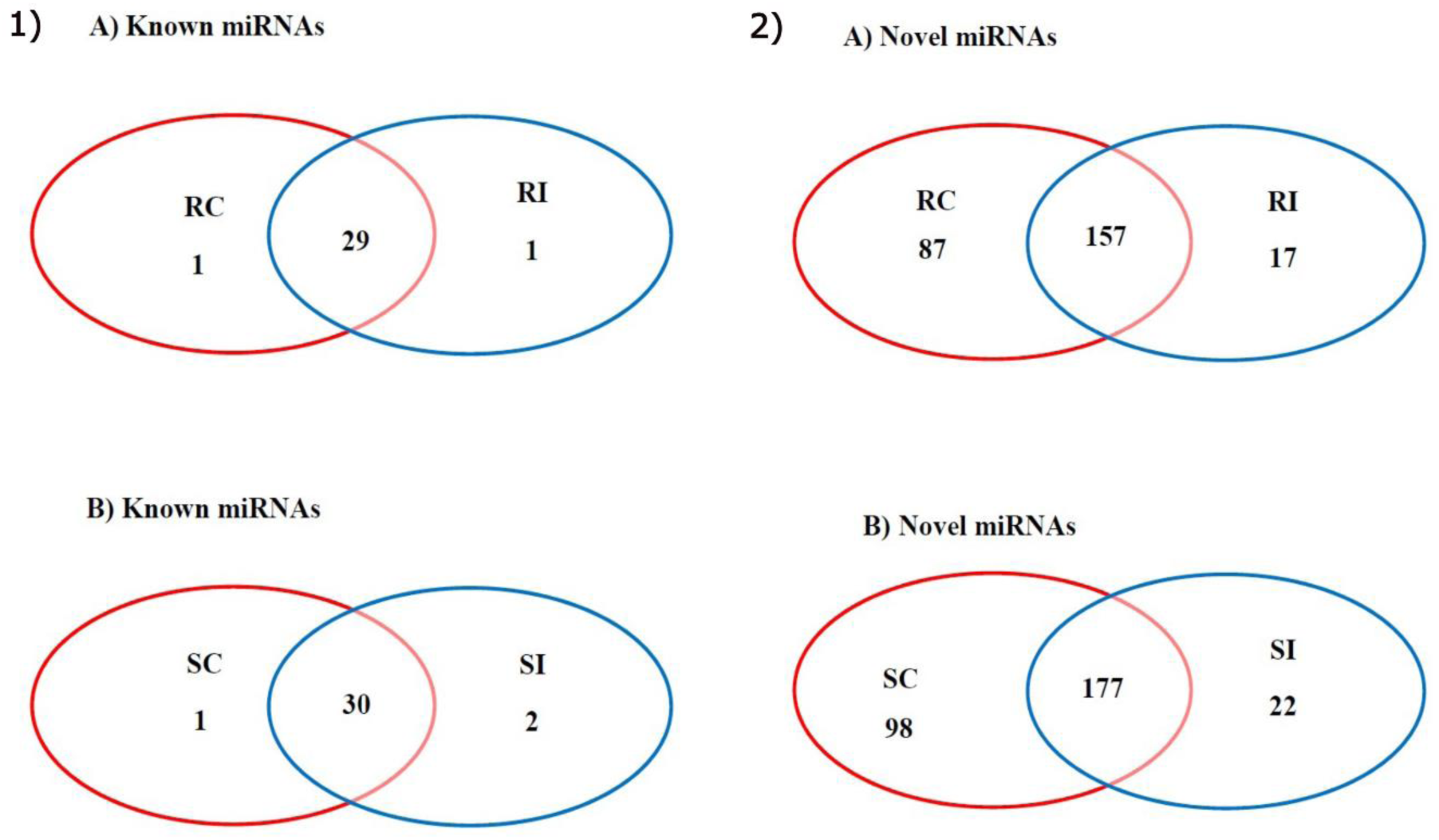
Venn diagram showing common and exclusively expressed miRNAs in resistance and susceptible chilli pepper. The red colour circle represents the control samples, whereas the blue colour circle represents the *P. capsici* infected leaf samples.1) known miRNAs in A) Resistance Control Vs. Resistance Infected and B) Susceptible inoculated; 2) Novel miRNAs in A) Resistance Control Vs. Resistance Infected and B) Susceptible Control Vs. Susceptible Infected.

### Identification and differential regulation of target genes

The result shows that most of the differentially expressed miRNAs-regulating target genes associated with *P. capsici* infection in SC vs SI leaf samples were novel miRNAs (Supplementary Table S5). To know the corresponding target genes of each of the miRNA during *C. annum – P. Capsic*i interaction, extensive target prediction was performed that resulted into identification of a total of 22,895 potential target genes, which was shown to be regulated by 79 known miRNA (corresponding to 24 miRNAs families) and 477 novel miRNAs in all four-leaf samples (Supplementary Table S6). Among these, 82.70 % (18,935) of the targets genes were shown as negatively regulated by miRNA-directed cleavage mechanism, and miRNAs translationally repressed 17.29 % (3,960) target genes. Potential miRNAs targets were predicted from our *C. annuum* transcriptome sequences (deposited in NCBI database under accession number RC: SAMN16251280; RI: SAMN16251798; SC: SAMN16251797; SI: SAMN16251799 on SRA as mentioned above. Hence, target gene annotation analysis revealed that these target genes were involved in different biological functions, including metabolic, cellular, biological processes, molecular activities, development process, resistance to biotic and abiotic stress response and response in the stimulus. Among these target genes, 30 genes were associated with defence response against *P. capsici* infection, which was regulated by 34 miRNAs, i.e. thirteen known and 21 novel miRNAs (Supplementary Table S5). The expression of few essential defence related target genes such as EP3-like gene, PLBRP, PDRP, PGUI, peroxidase 42, PRTA (PTI6), AP2-like ERTF(TOE3) and defensin-like protein genes was regulated by Pz-novel-miR1028-1, Pz-novel-miR428-1, Pz-known-miR2059-1, Pz-known-miR803-1, Pz-known-miR1872-1, Pz-novel-miR160-1, Pz-known-miR2560-1, and Pz-novel-miR204-1, respectively. The pathway for chitin biosynthesis (prominent cell wall component of fungus) was observed to be negatively regulated by Pz-novel-miR1028-1 by cleaving the target NP_001311656.1endochitinase EP3-like precursor gene in the SI leaf sample (Fig. 6B). Compared to RI, the expression of NP_001311656.1endochitinase EP3-like precursor gene was down-regulated in SI, suggesting its vital role in modulating the biosynthesis of chitin (Fig. 6B).

### Validation and expression profiling of selected miRNAs and their target genes

To further validate and check the expression pattern of identified miRNAs, four known (Pz-known-miR803-1, Pz-known-miR2059-1, Pz-known-miR2560-1 and Pz-known-miR2560-1) and four novel miRNAs (Pz-novel-miR428-1, Pz-novel-miR1028-1, Pz-novel-miR160-1, and Pz-novel-miR204-1) targeting eight corresponding key genes (*PGUI, PDRP, AP2-ERTF, POD, PLBRP, EP3, PRTA-PT16 like gene, DLP*) associated with defence response during *P. capsici* infection were selected. RT-qPCR and RA-PCR were employed for experimental validation of these miRNAs and their targets, respectively, in contrasting chilli pepper landraces. The amplified products were resolved on 2% agarose gel, and amplicons of respective miRNAs and their target mRNAs were detected. The RT-qPCR data was found consistent with the small RNA sequencing data in both resistant and susceptible landraces. The RT-qPCR analysis reveal that, the Pz-novel-miR428-1, Pz-known-miR803-1, Pz-novel-miR1028-1, Pz-novel-miR160-1, Pz-known-miR2059-1, Pz-known-miR2560-1, and Pz-novel-miR204-1 were differentially up-regulated while Pz-known-miR1872-1 was down-regulated in RC vs RI leaf (Fig 6 A-D and 7 A-D). However, all of the eight analyzed miRNAs were differentially regulated in SC vs SI leaf samples (Fig 6 A-D & 7 A-D). Compared to RC vs RI, the expression pattern indicated a varying accumulation of all the studied miRNAs with the highest differential regulation in SC vs SI (Fig. 6 and 7). Amongst all miRNA-target pair, the Pz-known-miR1872-PODs pair showed perfect inverse relation in all four samples (Fig 7D). Fig. 5 depicts the relative positions of the three primer sets of each target genes for RA-PCR. The 5′, mid, and 3′ regions of each target gene signify upstream, target site and downstream site from the potential cleavage site, respectively. Surprisingly, all the miRNAs’ expression at least in one of the four samples was negatively correlated with their target transcripts for the RA-Mid region (Fig. 6 & 7). Further, the RA-Mid region’s accumulation pattern of nearly all the transcripts was found to be less than the RA 3′ region (Fig. 6 & 7), suggesting that these target genes are cleaved by corresponding miRNAs.

**Fig. 5.**
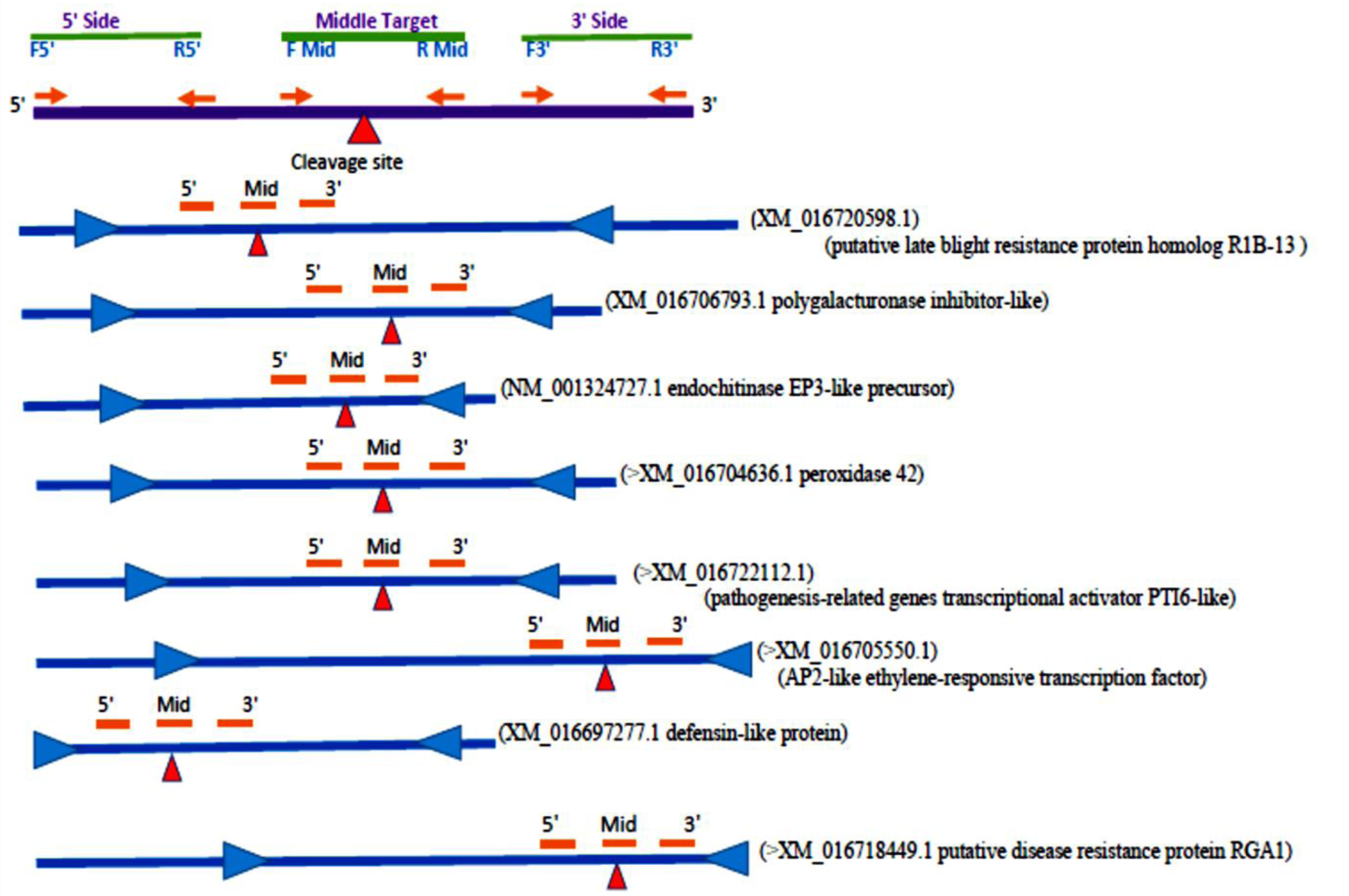
The diagram of regional amplification of qRT-PCR of target genes: A) The three site of primers used for regional amplification of target gene cleaved by miRNAs (5’ side both pair of forward and reverse primers, middle side both forward and reverse primers, middle side both forward and reverse primers, 3’ side both pair of forward and reverse primers); B) The miRNA cleavage site 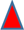) of eight target genes associated to the infection of *P. capsici*, the start and stop codon was illustrated by (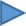 and 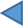) respectively.

**Fig. 6.**
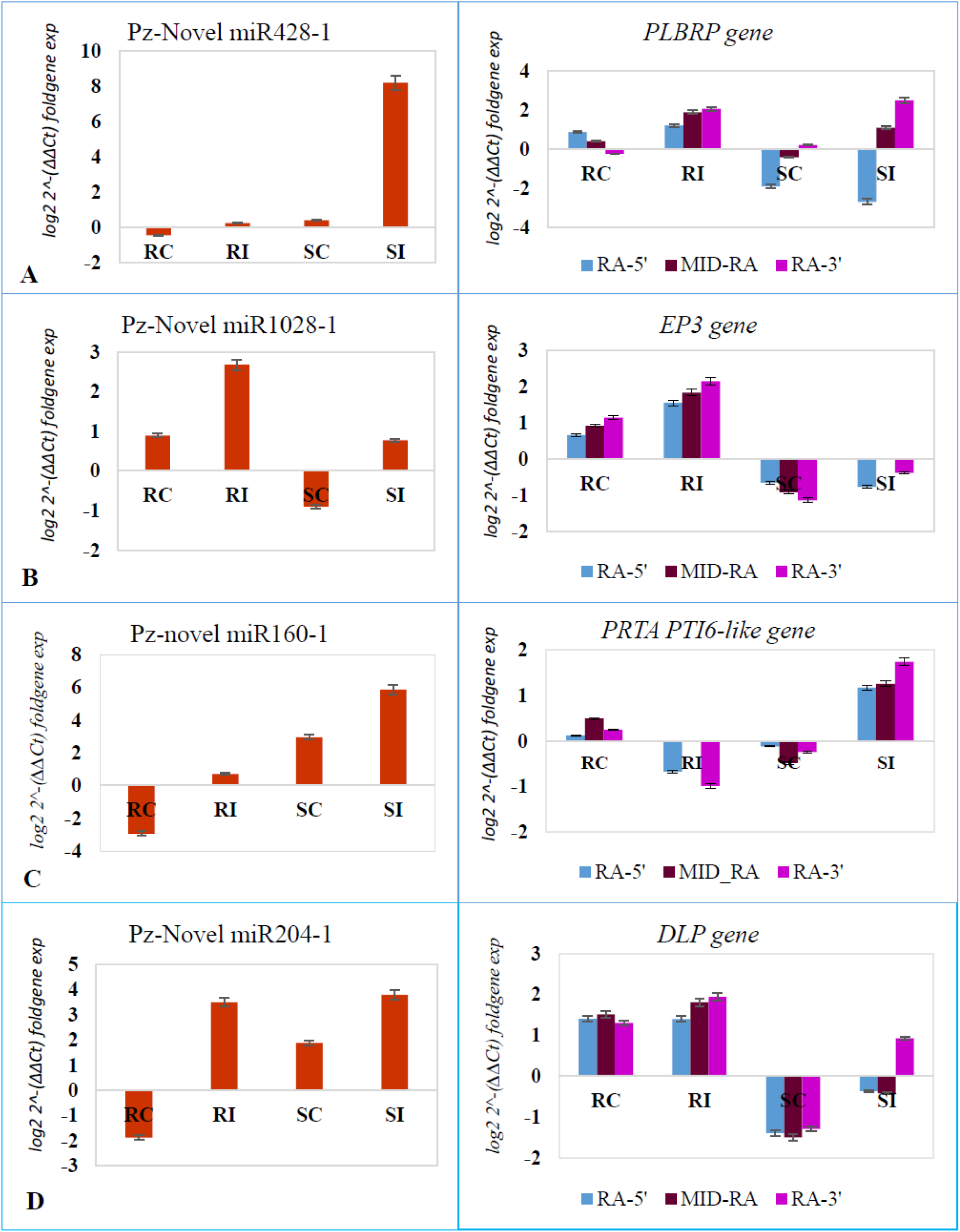
The RT-qPCR result analysis of the randomly selected DEGs of miRNAs regulating its corresponding identified target gene associated with defence response against *P. capsici* infection: A) Pz-novel-miR428-1 with its target putative late blight resistance protein homolog (PLBRP) RIP-13 gene, B) Pz-novel-miR1028-1 gene with its target endochitinase (EP) EP3-like precursor gene, C) Pz-novel-miR160-1 regulating its target PRTA-PTI6-like gene, and D) Pz-novel miR204-1 gene-regulating its target defensin-like protein(DLP) gene.

**Fig. 7.**
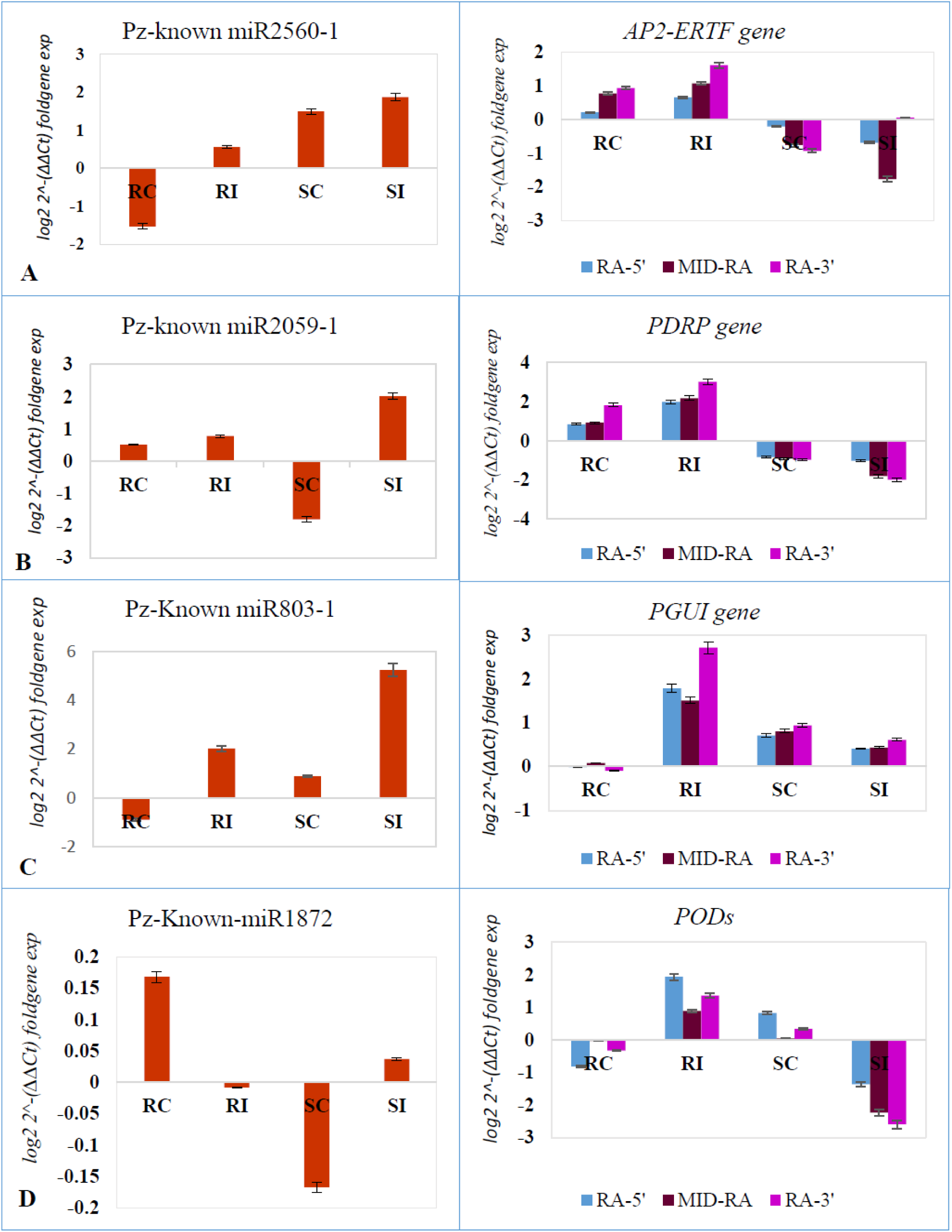
The RT-qPCR result analysis of the randomly selected DEGs of miRNAs regulating its corresponding identified target gene associated with defence response against *P. capsici* infection: A) Pz-known-miR2560-1(Can-miR172) gene-regulating its target AP2-like ethylene-responsive transcription factor (AP2-ERTF) TOE3 gene, B) Pz-known-miR2059-1 gene-regulating its target putative disease resistance protein (PDRP) RGA1 gene, C) Pz-known-miR803-1 with its target polygalacturonase inhibitor like (PGUI) gene, and D) Pz-known-miR1872-1(Can-miR397) with its target peroxidase 42 genes (PODs).

## Discussion

Owing to chilli pepper’s significant economic importance (*C. annuum* L.), deciphering the crucial role of miRNAs in modulating disease response is essentially required. As discussed in the introduction section, that despite numerous reports on miRNAs in chilli pepper, the molecular mechanism underlying *C. annuum-P. capsici* pathosystem is still a virgin field. Therefore, we performed a three-step experiment, i.e. (1) genome-wide identification of known and novel miRNAs in response to *P. capsici* infection in resistant and susceptible chilli pepper landrace; (2) identification of their corresponding target genes; and (3) validation and expression profiling of vital miRNA-target pairs associated with defence response in two contrasting chilli pepper landraces (resistant: GojamMecha_9086; susceptible: Dabat_80045).

The advancement of NGS technology coupled with bioinformatics analysis has revolutionized the identification of several families of defence-related known and novel miRNAs in plants exposed to various fungal pathogen. Based on the miRNA sequence analysis, 79 known conserved miRNAs corresponding to 24 miRNA families and 477 novel miRNAs were identified in the four chilli pepper libraries upon infection with *P. capsici* (Table 3). Although there is no report on miRNAs in chilli pepper during *P. capsici* infection, however, the current small RNA sequencing analysis obtained a large number of novel miRNAs (477 novel miRNAs) in four libraries as compared to the 35 novel miRNAs identified in ten libraries of *C. annuum,* which are involved in disease resistance-related genes such as verticillium wilt disease resistance proteins, late blight resistance proteins and NBS-LRR type disease resistance proteins (Shin *et al*., 2013). Tao and co-workers (Tao Li *et al*., 2015) were identified 71 miRNA families in two genotypes of *C. annuum* (i.e. tolerant hot pepper cultivar ‘R597’ (CaR) and the sensitive cultivar ‘S590’ (CaS) under high temperature and high air humidity stress. Din and co-workers (Din *et al*., 2016) identified and characterized a total of 88 new miRNAs (from 81 miRNA families) in chilli using expressed sequence tags (ESTs). Furthermore, a total of 59 known miRNAs and 310 novel miRNAs were identified from four libraries involved in fruit development and quality in *C. annuum* (Zou *et al*., 2017). Also, Robles and co-workers (Robles *et al*., 2019) identified 104 conserved miRNAs belonging to 37 families and 61 novel in miRNAs in tomato under five biotic and abiotic stress conditions (drought, heat, *P. syringae* infection, *B. cinerea* infection, and herbivore insect attack with *Leptinotarsa decemlineata* larvae). Comparatively, the known conserved miRNAs were highly expressed than the novel miRNAs in the treated sample of resistant and susceptible chilli pepper landrace. As identified, conserved miRNAs vary significantly among families (Cao et al., 2014); we also observed the same trends of variation of many miRNAs (12) for the can-miR169 family followed by can-miR395 and can-miR172 families. In line with previous reports of Cao *et al*. (2014) and Zhang *et al*. (2018), we also observed more significant variation in the abundance of clean read length distribution of mature miRNAs ranging from 15-29 nt. across all the libraries with 24 nt. being the most abundant (Fig. S2). According to Han *et al*. (2013), most small RNAs were found to be in the range of 20-24nt size, with the 24 nt. class being the most represented of the non-redundant species (75.83 %), followed by the 23 nt. (7.31 %) in *Caragana intermedia* by high-throughput sequencing under salt stress.

**Table 3.** The number of differentially expressed miRNA in control and infected leaf sample library combination of RC vs. RI and SC vs. SI.

| Differential combination<br>(Control_vs_ treated) | Type of miRNA | Number of down-regulated miRNA | Number of up-regulated miRNA |
| --- | --- | --- | --- |
| RC_vs_RI | Known | 4 | 25 |
|  | Novel | 6 | 151 |
| SC_vs_SI | Known | 5 | 25 |
|  | Novel | 32 | 144 |

As shown in Table 2, compared to known miRNAs, we observed differential expression of a more significant number of novel miRNAs in SC vs SI with nearly 32 novel miRNAs being down-regulated; supporting the report of Shin and co-workers (Shin *et al*., 2013), reported that some of the novel miRNAs were expressed at a high level in hot Pepper (*C. annuum*). However, Zou and co-workers (Zou *et al*., 2017) had reported a low expression level of novel miRNAs associated with fruit development and quality in *C. annuum*. This pattern of down-regulation of a large number of novel miRNAs in susceptible, infected chilli pepper landrace might signify miRNA depended on response in modulating disease susceptibility in comparison to resistant landrace by negatively regulating several target genes possibly associated with *P. capsici* infection and intensifying the vulnerability of the treated susceptible genotype to the pathogen infection. Further, differential expression of a nearly equal number of known miRNAs in both the landrace could have a nearly equal contribution of miRNA dependent disease response modulation. Further work on characterizing these novel miRNAs’ role using a reverse genetic approach shall impart more understanding of both the landrace’s resistance and susceptibility mechanism.

Enrichment analysis of identified known and novel miRNAs (∼34) showed their possible involvement in regulating target genes (∼30) associated with resistance against *P. capsicum* infection. Compared to SI, conserved miRNA families such as miR482, miR159, and miR166 were the most abundantly expressed and exhibited a high read count 156540.84, 211794.66, and 41876.76, respectively, in RI. This result agrees with previous reports in *C. annuum* and *S. lycopersicum,* where these conserved miRNAs were relatively less expressed than our result*s (*Shin *et al*., 2013; Robles *et al*., 2019). This shows that these conserved miRNAs have a more specific role in *C. annuum* in response to *P. capsici* infection. Despite no previous report documented on the role of miR166 in *C. annuum*, there was reported the role of miR166 in fungal pathogen infection in other plants. For instance, activated *mIR166k-166h* expression in rice plant showed enhanced infection resistance by the fungal pathogens *M. oryzae* and *F. fujikuroi,* in which the *ethylene-insensitive 2* (*EIN2*) gene was identified as a novel target gene for miR166k (Segundo *et al*., 2018). Satendra and co-worker (Satendra *et al*., 2020) identified an induced expression of miR166 under *R. solani* in susceptible and resistant rice cultivars, suggesting these may be basal response regulators against the *R. solani* pathogen. Furthermore, an increase in the accumulation of miR166 and miR159 in cotton plant in response to fungal pathogen *V. dahliae* infection reported (Jin and Guo, 2018). Nevertheless, *Pz-novel-miR1981-1* was the most abundantly expressed with TPM value of 21,280.07 and 15,007.33 in RI and SI samples, respectively, signifying their pathogen and genotypes dependent response to modulate the disease tolerance, which needs further functional investigation to ascertain the regulatory role during *P. capsici* infection.

Target genes identification revealed that most of the identified miRNAs were shown to regulate more target genes associated with resistance against *P. capsici* infection. The Pz-novel-miR3509-1 was observed to have 186 target genes, of which, six target genes (putative disease resistance protein At1g50180 isoform X2, late blight resistance protein R1-A-like, putative late blight resistance protein homolog R1B-14, disease resistance RPP13-like protein 4, ethylene-responsive transcription factor ERF071-like, and putative late blight resistance protein homolog R1A-10) were associated with defence response against *P. capsici* infection. Similarly, Pz-known-miR2059-1(mir172) regulated 141 target genes, including two (putative disease resistance protein (PDRP) RGA1 and leucine-rich repeat receptor-like tyrosine-protein kinase PXC3) genes associated with disease modulation. The RT-qPCR analysis revealed, high expression and differential up-regulation of conserved miRNAs, i.e. Pz-known-miR2059-1(can-mir172), Pz-known-miR2560-1(can-miR172), and Pz-known-miR1872-1(can-miR397) were exhibited inversely regulating their corresponding target putative disease resistance protein (PDRP) RGA1 gene, AP2-like ethylene-responsive transcription factor (AP2-ERTF) TOE3 gene, and peroxidase 42 genes (PODs) expression associated with *P. capsici* infection in SI landrace (Fig. 6). However, comparatively the expression of Pz-known-miR1872-1(can-miR397), Pz-known-miR2059-1(can-mir172), Pz-known-miR2560-1(can-miR172), and Pz-known-miR803-1(can-miR399) were shown to be low in RI landrace, indicating higher expression of their corresponding target genes i.e. peroxidase 42 genes (PODs), putative disease resistance protein *(PDRP) RGA1 gene, AP2-like ethylene-responsive transcription factor (AP2-ERTF) TOE3 gene, and polygalacturonase inhibitor like (PGUI) gene*, respectively, which might be one of the reasons of increased resistance to *P. capsici* infection in GojamMecha_9086. Wang *et al*. (2018), up-regulation of several conserved miRNAs including miR159, miR168, miR169, miR172, miR393, miR398, and miR396 in *S. lycopersicum* and *S. habrochaites* during *B. tabaci* infestations suggesting that conserved miRNAs play essential roles in defence. Our results indicated that leaf rust disease-resistance locus receptor-like protein kinase, one of the targets of can-miRNA398, was negatively regulated in *C. annuum* under the *P. capsici* treatment with more significant accumulation in RI compared to SI, signifying its vital role in modulating disease resistance more significantly in resistant chilli pepper. The expression difference in can-miRNA398 - leaf rust disease-resistance locus receptor-like protein kinase target pair might lead to variation in disease resistance response between resistant and susceptible genotype against *P. capsici* infection. The miRNA398 was negatively regulated pathogen-associated molecular patterns (PAMPs) induced callose deposition and disease resistance to bacteria, and the overexpression of miR398b in plants showed enhanced susceptibility to both virulent and non-pathogenic strains of *P. syringae*, indicating its essential role in disease resistance (Zhou *et al.,* 2010).

Among the total predicted target genes, 82.70% (18935) of the targets genes were negatively regulated by miRNA-directed cleavage mechanism, while miRNAs transitionally repressed 17.29%(3960) target genes. The current result shows an inverse correlation between the expression profiles miRNAs and their target genes which is in agreement with the previous report in *S. lycopersicum* during biotic and abiotic stress (Robles *et al*., 2019). In the current study, from 22,895 identified target genes, 30 targets genes were directly associated with resistance mechanism against *P. capsici* infection, which was regulated by eleven mature known and 21 novel miRNAs. Robles et al. (2019) and Zhang et al. (2018) reported that the expression of many single target genes was observed to be regulated by different miRNAs of different families during *C. capsici* infection in chilli pepper. For instance, the putative late blight resistance protein homolog R1B-17 gene was regulated by three miRNAs, i.e. Pz-known-miR1099-1(can-miR159), Pz-known-miR1382-1 (can-miR319), Pz-known miR2610-1 (can-miR319) and the polygalacturonase inhibitor-like genes were regulated by four miRNAs, i.e. Pz-known-miR803-1 (can-miR399), Pz-novel-miR1405-1, Pz-novel-miR3245-1, and Pz-novel-miR3502-1. The target squamosa promoter-binding-like protein 6(SBP) is regulated by five miRNAs, i.e. Pz-known-miR1186-1, Pz-known-miR3461-1, Pz-novel-miR227-1, Pz-novel-miR1107-1, and Pz-novel-miR2181-1. Gong and co-worker have reported the ivolvement of target SBP the defense response of pepper to *Phytophthora capsici* by regulating defense-related genes(Gong et al., 2020^b^). Moreover, our result shows, the Pz-known-miR2059-1(can-miR172) regulated expression of target AP2-like ethylene-responsive transcription factor TOE3 isoform X1 gene, which activates signalling pathway during *P. capsici* infection, and it was in agreement with the report on the role of miR172 in *S. lycopersicum* and *S. habrochaites* under *Bemisia tabaci* infestations (Wang *et al*., 2018). The RT-qPCR result shows the accumulation of Pz-known-miR1872-1(can-miR397) inversely targeting its corresponding target peroxidase 42 genes in SI than in RI landrace that possibly creates differential disease response between the resistance and susceptible genotype of chilli pepper. The peroxidase enzyme participated in lignin and suberin formation, crosslinking of cell walls components, and synthesis of phytoalexins, leading to the hypersensitive response (HR), a form of programmed host cell death at the infection site associated with limited pathogen development (Almagro *et al*., 2009). The PR-9 family of peroxidase enzyme have involved in cell wall reinforcement by catalyzing lignification, leading to enhanced resistance against multiple pathogens (Passardi *et al*., 2004).

Pz-novel-miR1028-1 and Pz-novel-miR177-1 regulate expression of gene encoding endochitinase protein involved in catalyzing the hydrolytic cleavage of the β-1,4 glycoside bond present in biopolymers of N-Acetyl-D-glucosamine, mainly in chitin the wall components of fungus (Ferreira *et al*., 2007). The CAChi2 chitinase is encoded by a single or two copy genes in the pepper genome (Kim *et al*., 2000). There was a report on the class IV chitinase, CaChitIV, from pepper plants (*C. annuum*), which interacted with the pepper receptor-like cytoplasmic protein kinase(CaPIK1) and promotes CaPIK1-triggered cell death and defence responses (Kim *et al*., 2015). NAC transcription factor expression 29-like targeted by Pz-novel-miR34-directed cleavage in chilli pepper has an analogous function with ath-miR164, and the decreased expression leads to increased expression of the NAC transcription factor in *A. thaliana*. The target NAC transcription factors are involved in regulating tissue development in response to biotic and abiotic stress, which was a positive regulator of cell death (Woo *et al*., 2009; Chuck and O’Connor, 2010). The expression of Pz-known-miR1186-1(∼8.0 fold) explicitly in SC leaf negatively was regulated the expression of target squamosa promoter-binding-like protein 6 (SBP) gene. However, expression of this target SBP in SI leaf enhaced susceptibility to *P.capsici* infection. *CaSBP11* has a negative regulatory effect on defense responses to *Phytophthora capsici* suggests their potentially significant role in plant defense(Gong *et al*., 2018; Gong *et al*., 2020^b^). Silencing of *CaSBP08* and *CaSBP11* enhanced resistance to *P. capsici* infection in *Nicotiana benthamiana* and pepper, respectively (Gong *et al*., 2020^a^; Gong *et al*., 2020^b^).

Moreover, among eight miRNAs validated, five miRNAs, i.e., Pz-novel-miR428-1, Pz-Novel-miR1028-1, Pz-known-miR1872-1, Pz-known-miR2059 and Pz-known-miR2560-1, negatively regulate the expression of their corresponding target PLBRP, EP3, PODs, PDRP gene and AP2-ERTF, respectively, in SI under *P. capsici* infection. However, the Pz-novel miRNA160-1, Pz-known miR803-1, and Pz-novel-miR204-1 were positively regulated with their corresponding target PRTA PTI6-like gene PGUI, and defensin-like protein gene, respectively, in SI leaf. Similarly, the expression of eight randomly selected miRNAs except, Pz-novel-miR160-1 positively regulate the expression of their corresponding target gene during *P. capsici* infection in the RI leaf sample. Our observation of differential expression of miRNA-target genes in both RI and SI coincides with several reports suggesting an inverse correlation between the miRNA and their corresponding target genes (Robles *et al*., 2019; Gupta *et al*., 2017).

In summary, the current small RNA sequencing of chilli pepper exposed to *P. capsici* infection in two contrasting, i.e. resistant and susceptible, chilli pepper landraces identified various conserved and novel miRNAs involved in regulating potential target genes associated with defence response against *P. capsici*. The differential expression behaviour of several conserved and novel miRNAs and their target genes in both RI and SI leaf samples of resistant and susceptible chilli pepper landrace suggests their paramount role in resistance mechanism under *C. annuum L. - P. capsici* pathosystem. The variation in the expression level of defence associated miRNAs and their corresponding target genes might be one reason for differential disease response between resistant and susceptible landraces under *C. annuum* L. infection. These results need further functional validation using reverse genetic and/or Crispr/cas9 approaches by selecting few defence associated miRNAs and their corresponding target genes to broaden our current understanding of miRNA based regulation of disease resistance during *C. annuum* L.-*P. capsici* pathosystem.

## Declarations

This research did not involve the use of any animal or human data or tissue.

## Author’s contributions

Conceptualization, Data curation, Formal analysis, Investigation, Methodology, Writing original draft, Software, Validation, T.R, Om.P.G and V.C., Project administration, Resource, V.C, Writing, Revising and Editing, Supervision, Om.P.G, and V.C.

**Declaration of competing interest**

## Acknowledgement

TR is gratefully acknowledging the Ministry of Science and Higher education, Ethiopia, for sponsoring a fellowship program. Besides, appreciation and many thanks go out to the Department of Bio and Nanotechnology, Guru Jambheshwar University of Science and Technology, Hisar, India, to prove all necessary laboratory facilities. Special thanks also go to all our lab mate who gave invaluable advice and feedback at lab work.

## Supplementary Fig

**Fig. S1.**
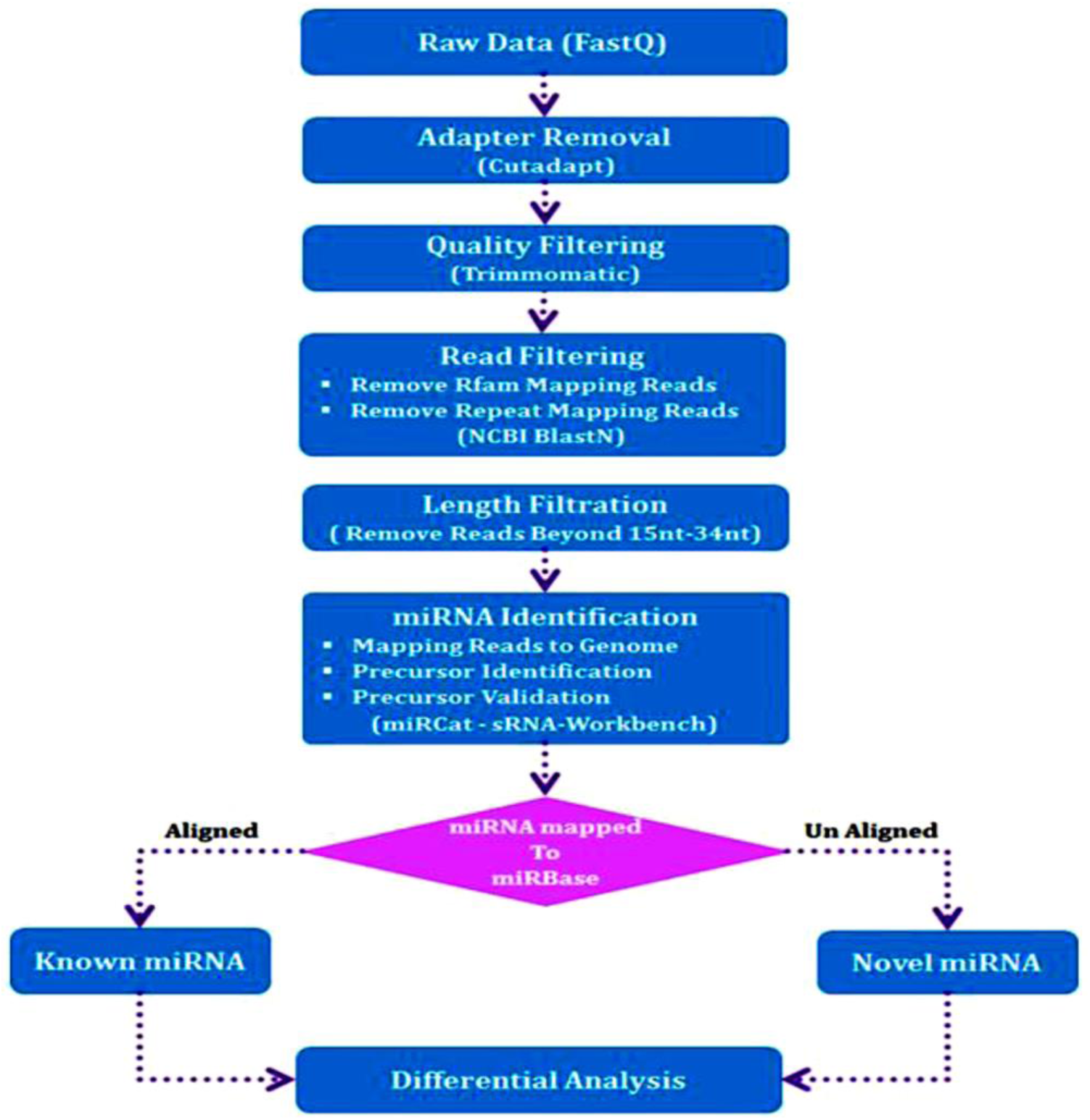
The bioinformatics workflow for small RNA sequencing and data analysis

**Fig. S2.**
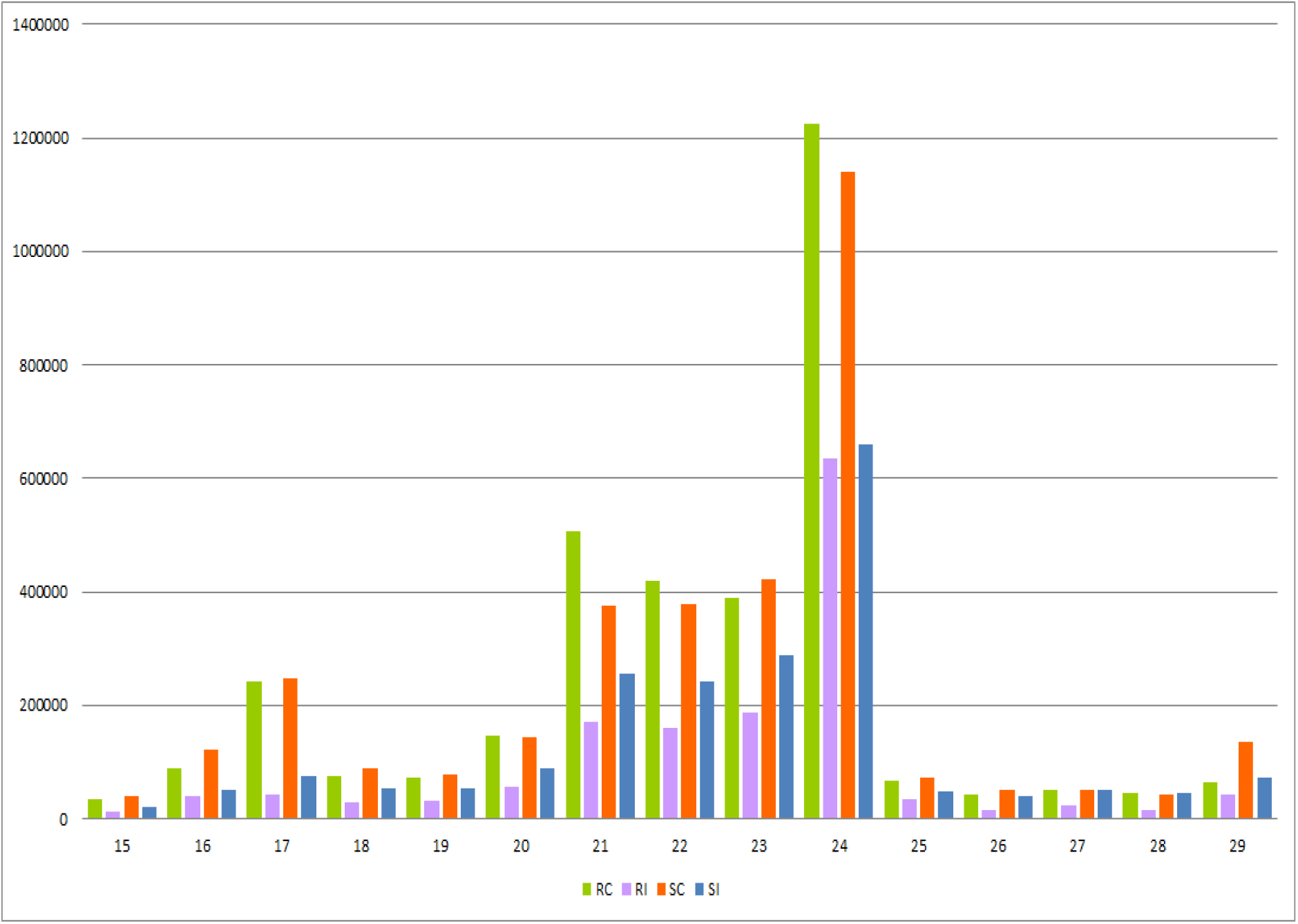
Length distribution of final cleaned reads (15-29nt.) for the four-leaf sample of chilli pepper

## Supplementary Tables

**Table S1. The designed primers for randomly selected differentially expressed miRNA genes for validation of miRNA Seq using RT-qPCR**

**Table S2.**
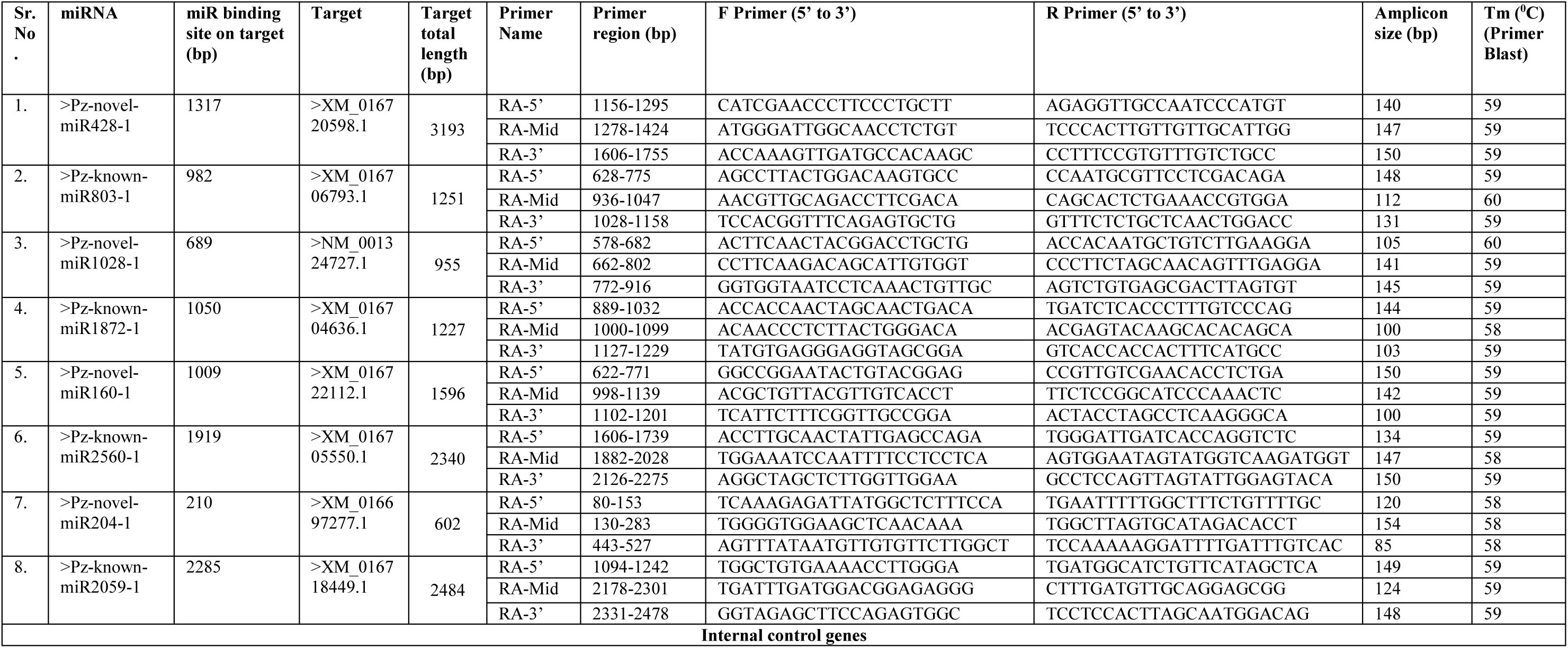
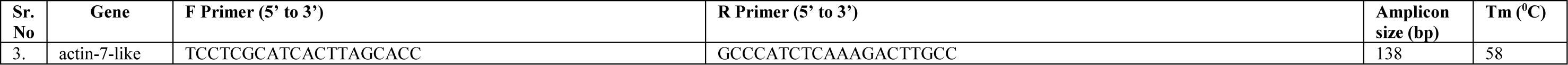
The designed primers for eight miRNAs target genes and exogenous actin-7-like primer randomly selected for validation using RT-qPCR

**Table S3.**
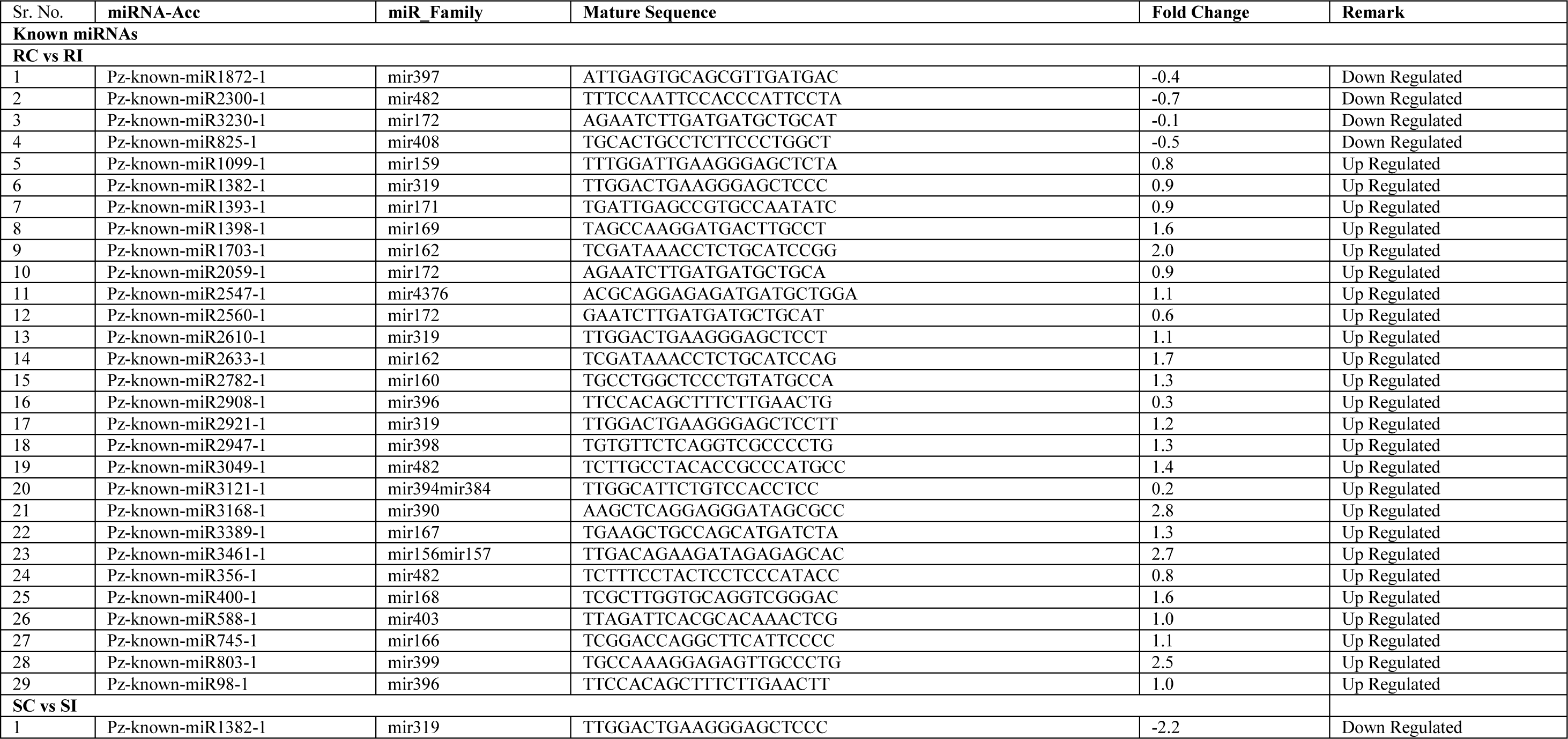
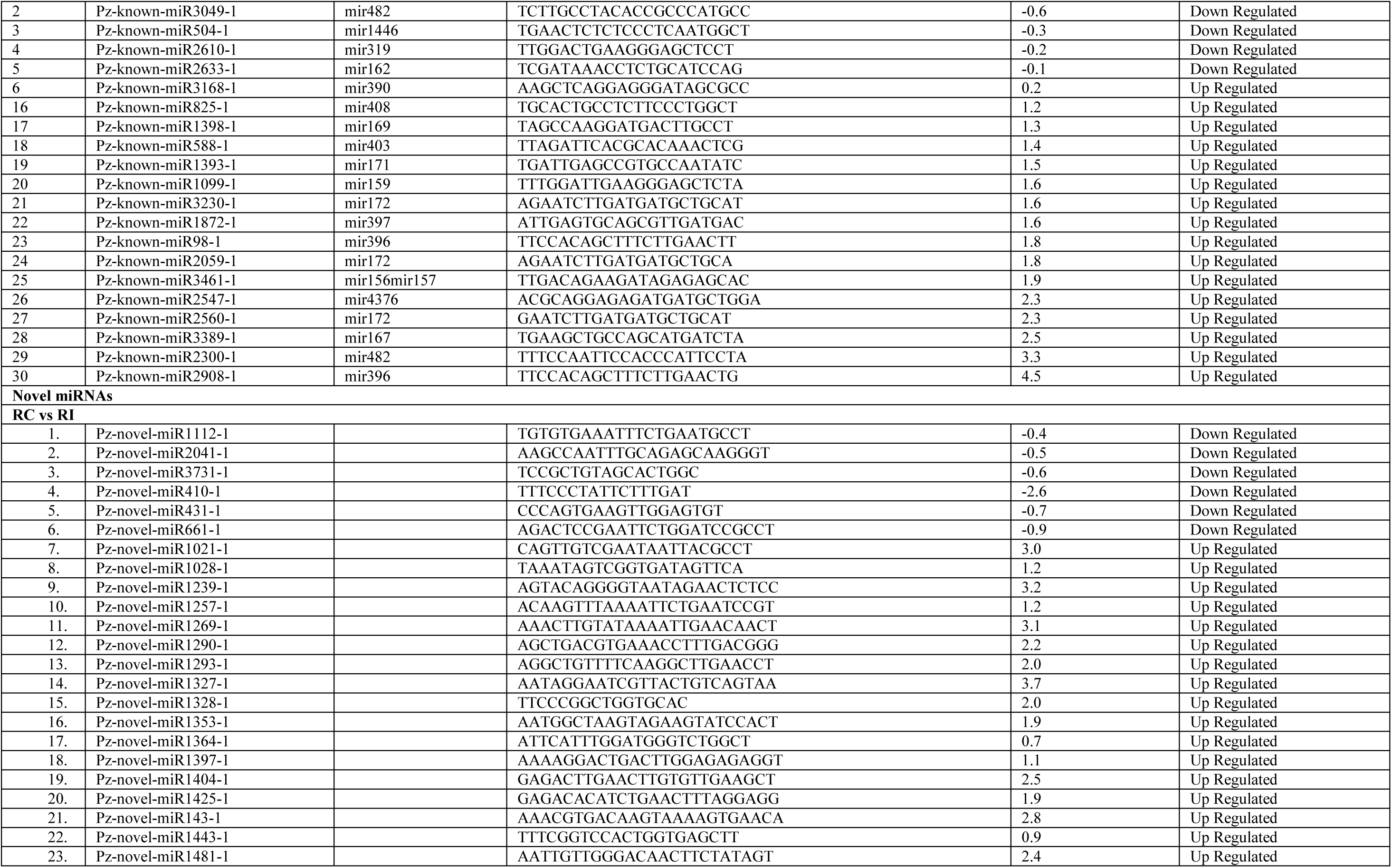
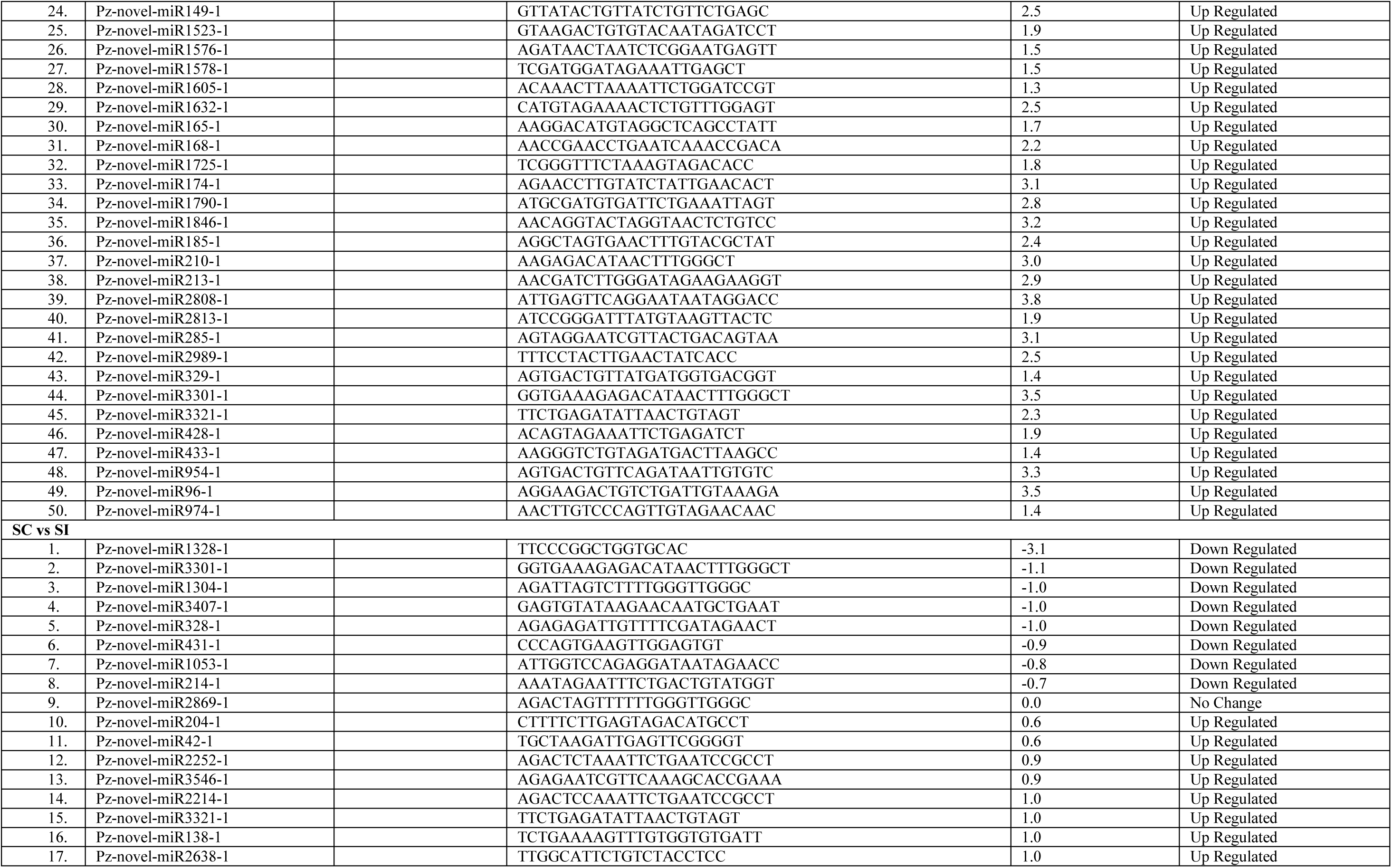
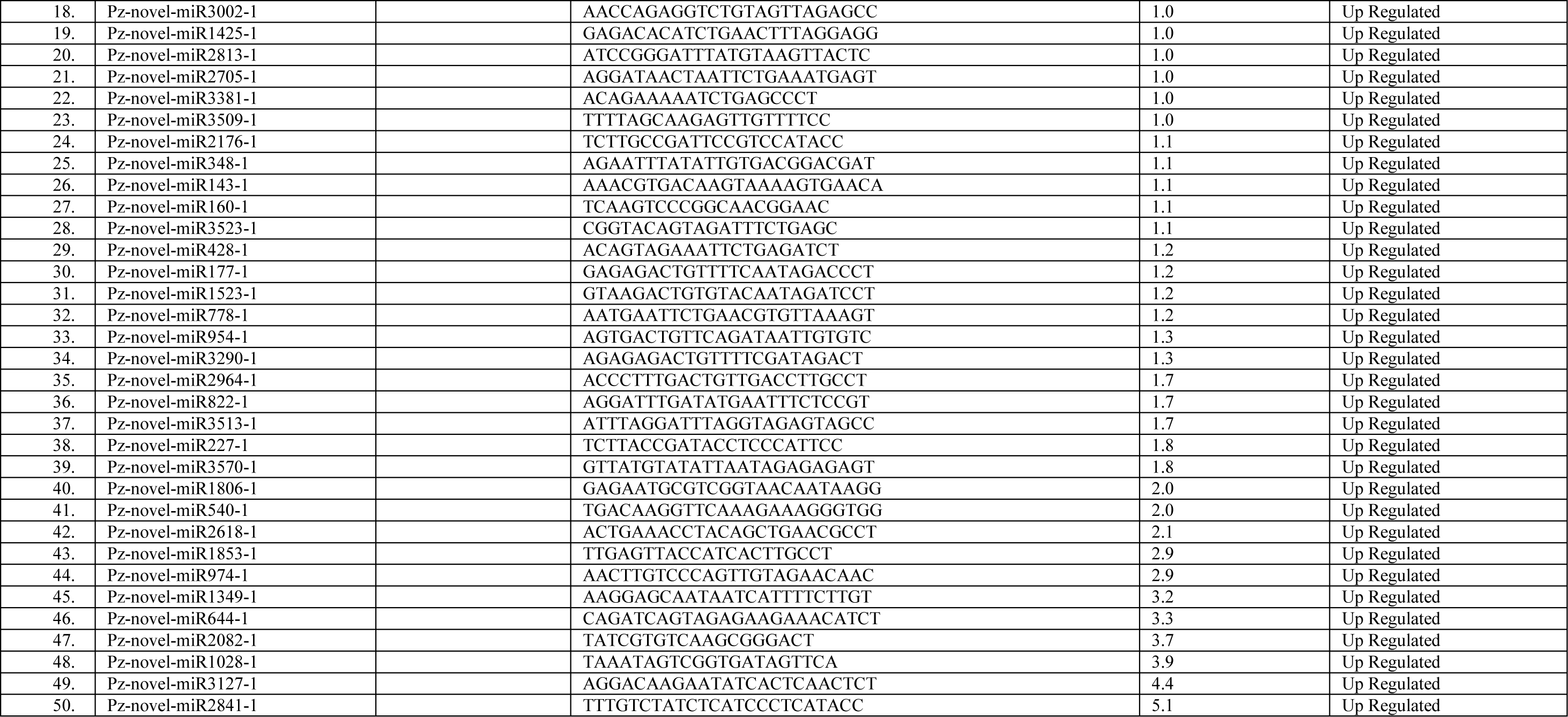
List of commonly expressed known and novel miRNA along with their fold change value in *C. annuum* L. under *P. capsici* infection.

**Table S4.**
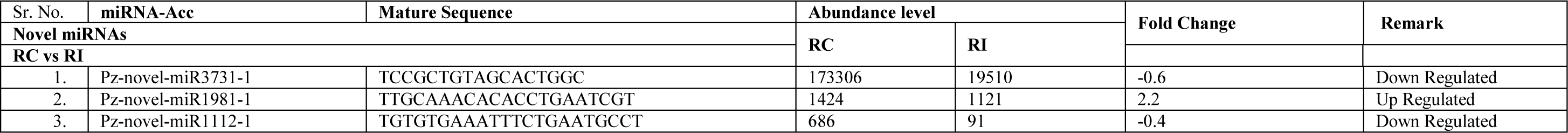
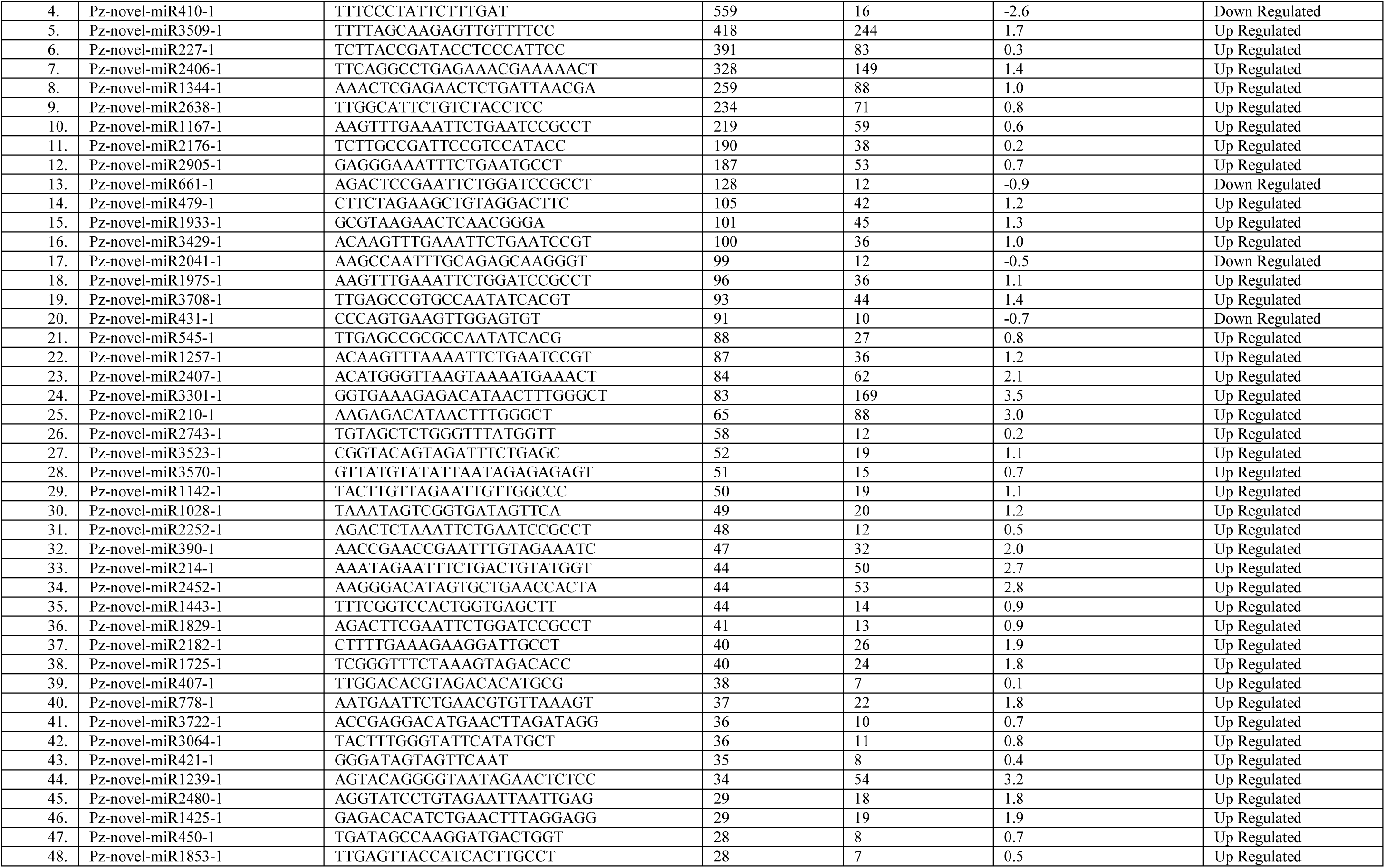
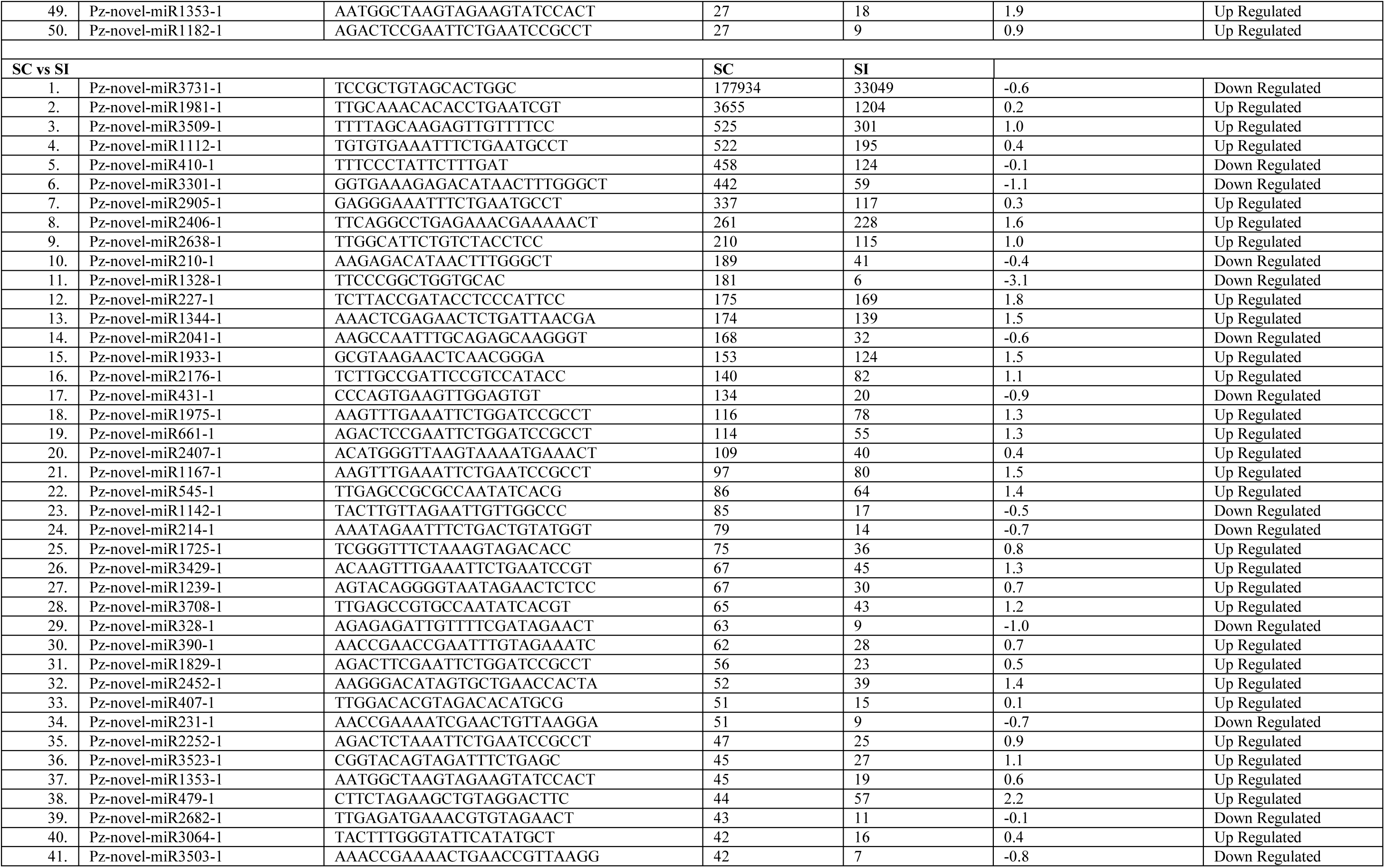
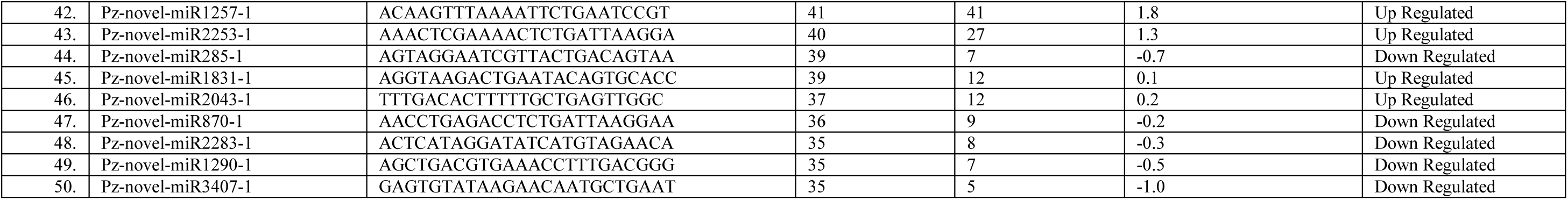
List of all the novel miRNA and their fold change value in RC vs RI and SC vs SI sample of *C. annuum* L. under *P. capsici* infection.

**Table S5.**
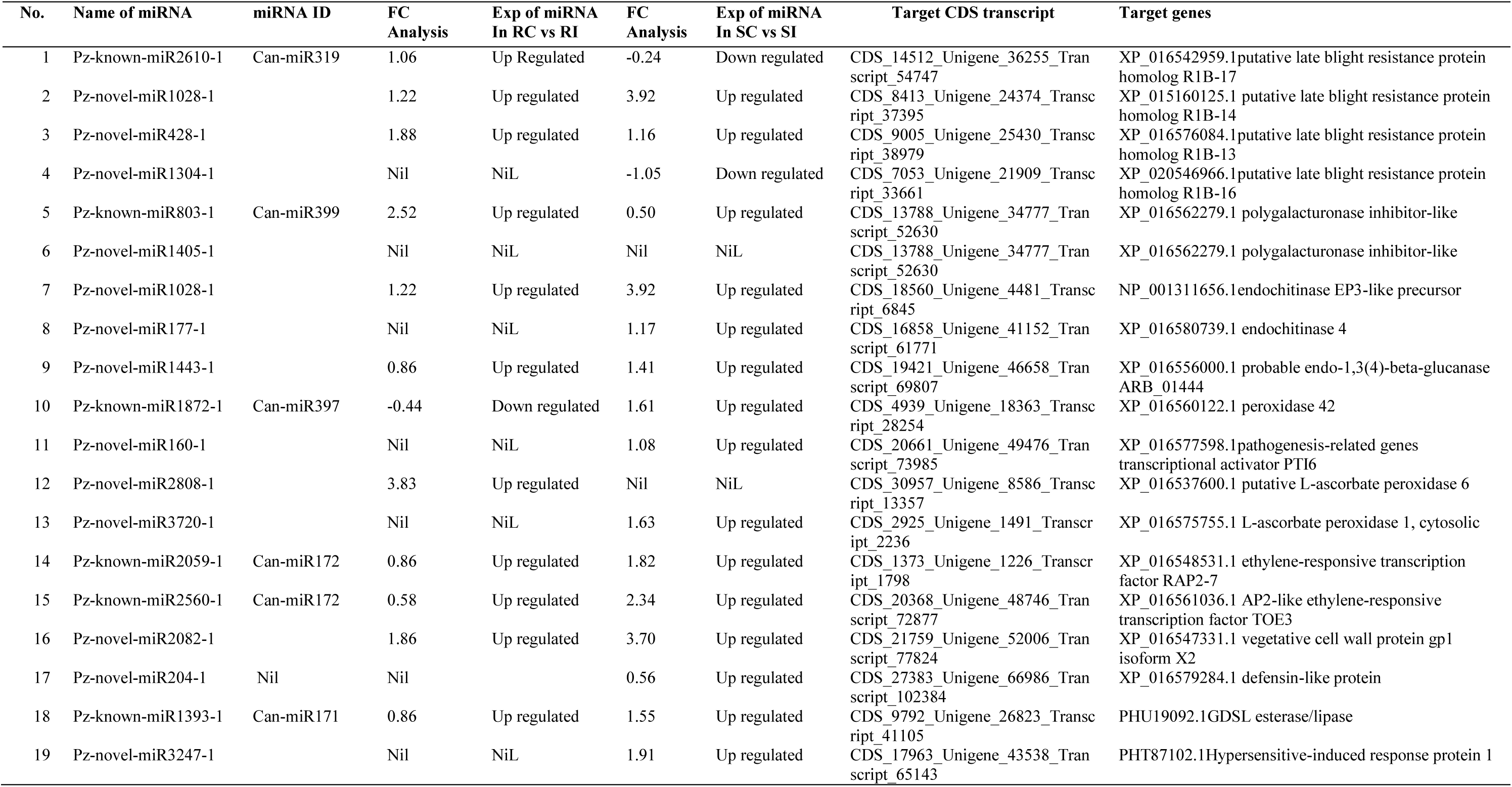
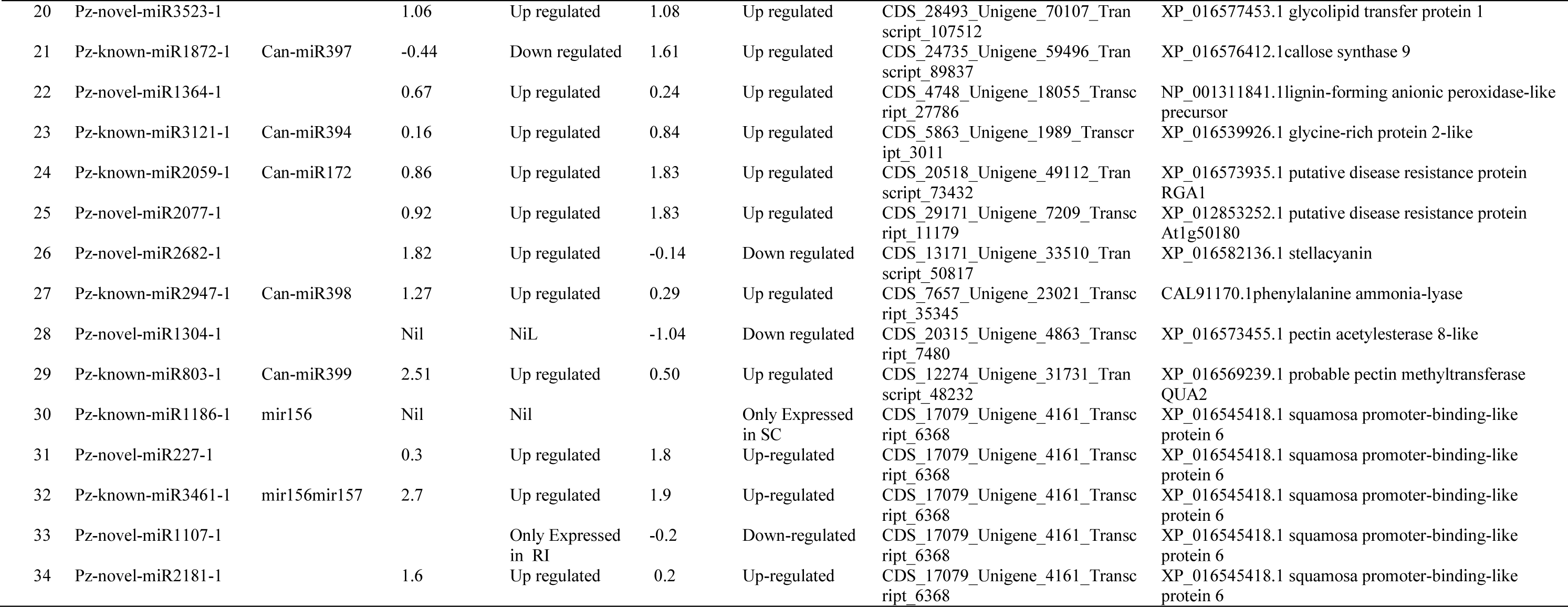
The differentially expressed genes of known and novel miRNA with their corresponding target genes associated with *P. capsici* infection in resistance and susceptible *C. annuum* L (in sample combination of resistance control and resistance inoculated and susceptible control and susceptible infected).

**Table S6.**
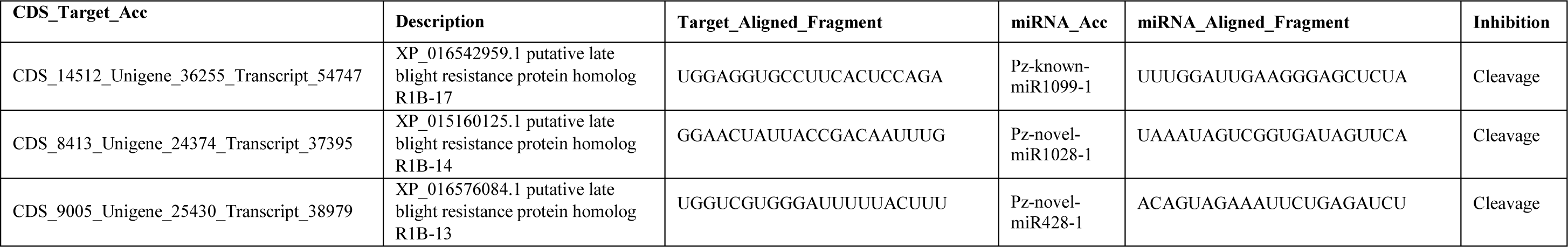
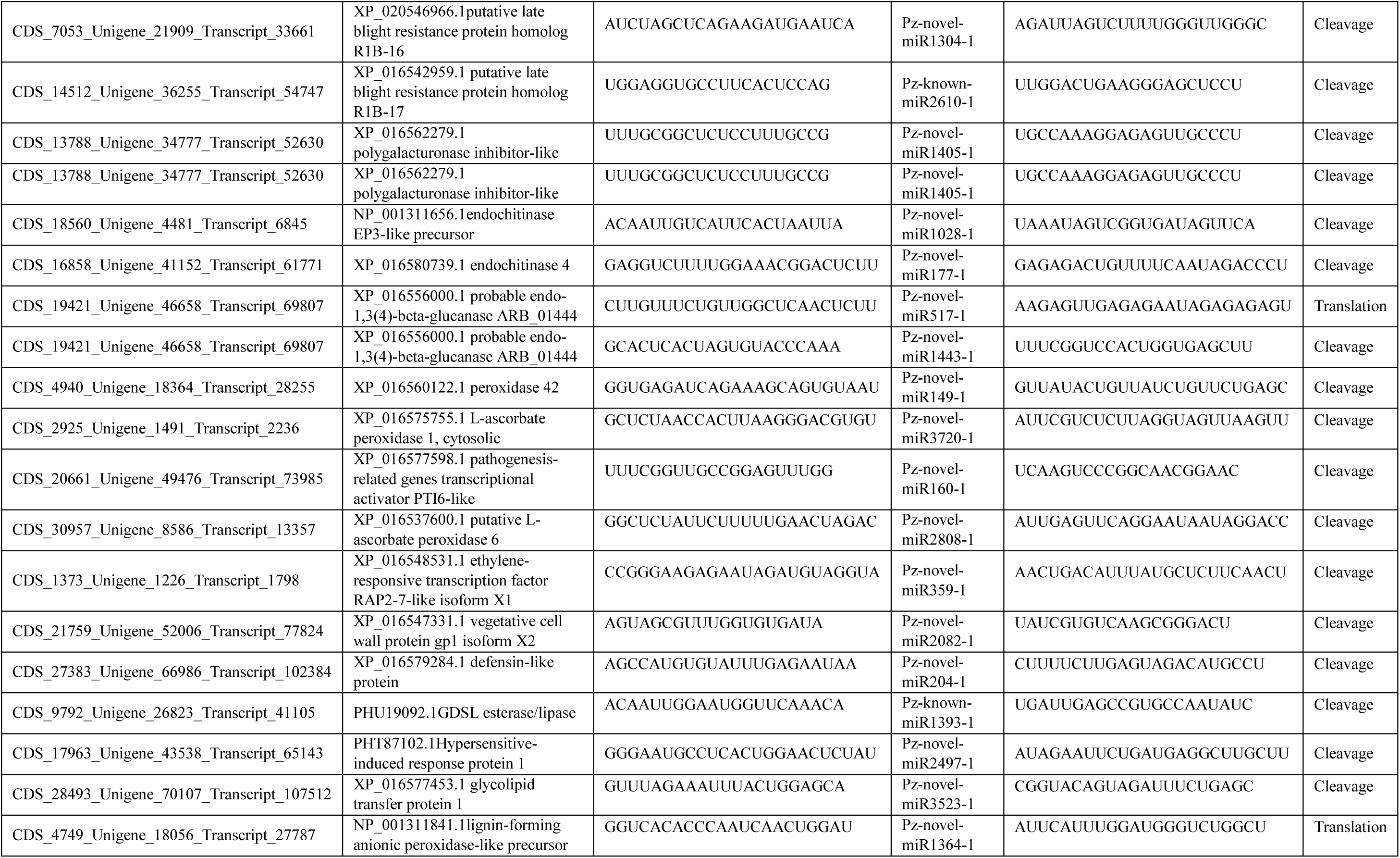
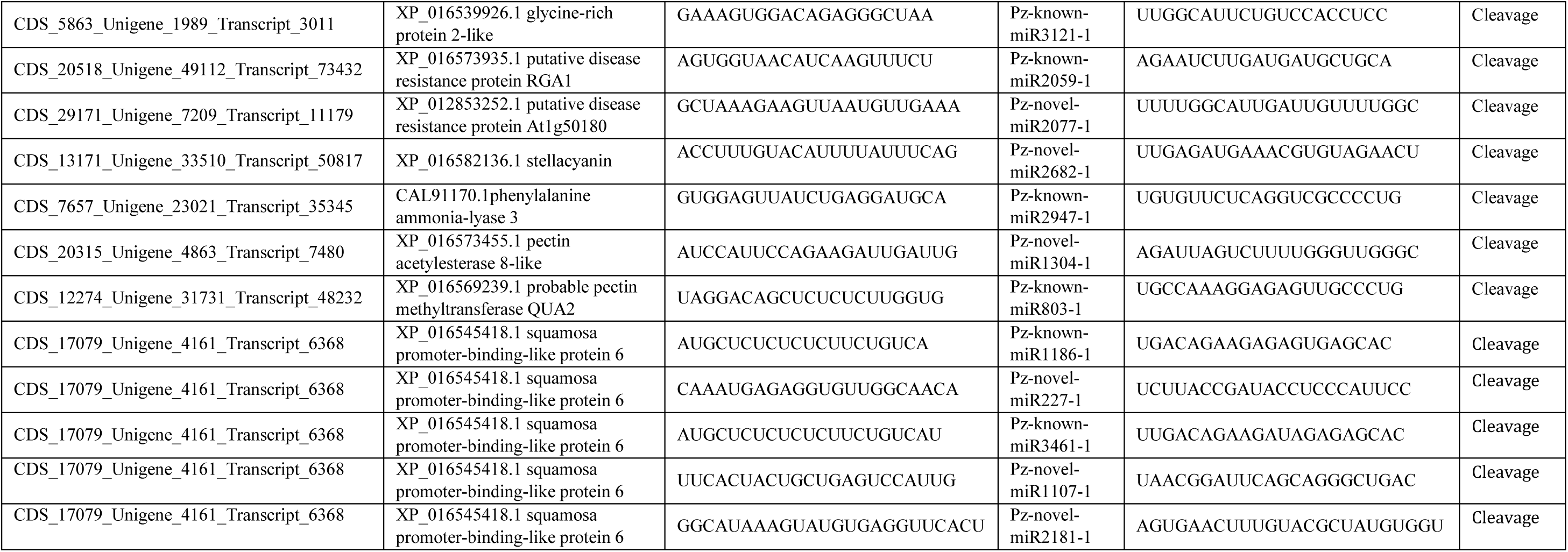
List of the target genes along with their potential corresponding miRNAs

## Reference

1. Bartel, D.P., 2004. MicroRNAs: Genomics, biogenesis, mechanism, and function. Cell, 116, 281–297.

2. Peng T, Sun H, Du Y, Zhang J, Li J, Liu Y, Zhao Q, et al., 2013. Characterization and Expression Patterns of microRNAs Involved in Rice Grain Filling. PLoS ONE 8(1): e54148. https://doi.org/10.1371/journal.pone.0054148

3. Ana Kozomara, Maria Birgaoanu, Sam Griffiths-Jones, 2019. miRBase: from microRNA sequences to function, Nucleic Acids Research, Volume 47(1), 155–162, https://doi.org/10.1093/nar/gky1141

4. Yin, Z., Li, Y., Han, X., & Shen, F., 2012. Genome-wide profiling of miRNAs and other small non-coding RNAs in the Verticillium dahliae-inoculated cotton roots. PloS one, 7(4), e35765. https://doi.org/10.1371/journal.pone.0035765

5. Chen L, Ren Y, Zhang Y, Xu J, Zhang Z, Wang Y., 2012. Genome-wide profiling of novel and conserved Populus microRNAs involved in pathogen stress response by deep sequencing. Planta, 235(5):873–883. doi:10.1007/s00425-011-1548-z

6. Gupta OP, Permar V, Koundal V, Singh UD, Praveen S.,2012. MicroRNA regulated defense responses in *Triticum aestivum* L. during Puccinia graminis f.sp. tritici infection. Mol Biol Rep.; 39(2):817–824. doi:10.1007/s11033-011-0803-5 https://doi.org/10.1007/BF00417941

7. Din, M., Barozai, M. Y. K., & Baloch, I. A., 2016. Profiling and annotation of microRNAs and their putative target genes in chilli (Capsicum annuum L.) using ESTs. Gene Reports, 5, 62–69. doi:10.1016/j.genrep.2016.08.010

8. Sha Yang, Zhuqing Zhang, Wenchao Chen, Xuefeng Li, Shudong Zhou, Chengliang Liang, Xin Li, Bozhi Yang, Xuexiao Zou, Feng Liu, Lijun Ou & Yanqing Ma, 2021. Integration of mRNA and miRNA profiling reveals the heterosis of three hybrid combinations of *Capsicum annuum* varieties, GM Crops & Food, 12:1, 224–241, doi:10.1080/21645698.2020.1852064

9. Zhang, H., Huang, S., Tan, J., Chen, X., & Zhang, M., 2020. MiRNAs profiling and degradome sequencing between the CMS-line N816S and its maintainer line Ning5m during anther development in pepper (Capsicum annuum L.). bioRxiv.

10. Hwang D-G, Park JH, Lim JY, Kim D, Choi Y, Kim S, Shin C, et al., 2013. The Hot Pepper (*Capsicum annuum*) MicroRNA Transcriptome Reveals Novel and Conserved Targets: A Foundation for Understanding MicroRNA Functional Roles in Hot Pepper. PLoS ONE, 8(5): e64238. https://doi.org/10.1371/journal.pone.0064238

11. Khalid Annum, Zhang Qingling, Yasir Muhammad, Li Feng, 2017. Small RNA Based Genetic Engineering for Plant Viral Resistance: Application in Crop Protection. Frontiers in Microbiology, 8(43). doi;10.3389/fmicb.2017.00043

12. Liu, S. R., Zhou, J. J., Hu, C. G., Wei, C. L., & Zhang, J. Z., 2017. MicroRNA-Mediated Gene Silencing in Plant Defense and Viral Counter-Defense. Frontiers in microbiology, 8, 1801. https://doi.org/10.3389/fmicb.2017.01801

13. Barrajon-Catalan, E., Alvarez-Martinez, F.J., Borras, F., Perez, D., Herrero, N., Ruiz, J.J., and Micol, 2020. Metabolomic analysis of the effects of a commercial complex biostimulant on pepper crops. Food Chem, 310:125818. doi:10.1016/j.foodchem.2019.125818. Epub 2019 Nov 7. PMID: 31787397.

14. El-Ghoraba AH, Javedb Q, Anjumb FM, Hamedc SF, Shaabana HA., 2013. Pakistani Bell Pepper (*Capsicum annum* L.): Chemical Compositions and its Antioxidant Activity. International Journal of Food Properties, 16(1):18–32. https://doi.org/10.1080/10942912.2010.513616

15. Chamikara, M. D. M., Dissanayake, D. R. R. P., Ishan, M., & Sooriyapathirana, S. D. S. S., 2016. Dietary, anticancer and medicinal properties of the phytochemicals in chili pepper (*Capsicum spp*.). Ceylon Journal of Science, 45(3), 5–20. doi: http://doi.org/10.4038/cjs.v45i3.7396

16. Babadoost, M. 2004. Phytophthora blight: A serious threat to cucurbit industries. <http://www.apsnet.org/online/feature/cucurbit/>.

17. Mao, W., J.A. Lewis, R.D. Lumsden and K.P. Hebbar, 1998. Biocontrol of selected soilborne disease of tomato and pepper plants. Crop Prod. 17: 535–542.

18. Authors: Kaori Ando, Sue Hammar, and Rebecca Grumet, 2009. Age-related Resistance of Diverse Cucurbit Fruit to Infection by *Phytophthora capsici*. J. AMER. SOC. HORT. SCI. 134(2):176–182. Doi.https://doi.org/10.21273/JASHS.134.2.176

19. T. Rabuma, O. P. Gupta, and V. Chhokar, 2020. Phenotypic characterization of chili pepper (*Capsicum annuum* L.) under *Phytophthora capsici* infection and analysis of genetic diversity among identified resistance accessions using SSR markers. Physiological and Molecular Plant Pathology. 112. https://doi.org/10.1016/j.pmpp.2020.101539

20. J.L. Andrés Ares, A. Rivera Martínez, J. Fernández Paz,2005. Resistance of pepper germplasm to Phytophthora capsici isolates collected in northwest Spain, Spanish *J*. Agric. Res. 3 (4) 429–436, https://doi.org/10.5424/sjar/2005034-170

21. Tiwari JK, Buckseth T, Zinta R, Saraswati A, Singh RK, Rawat S, et al., 2020. Genome-wide identification and characterization of microRNAs by small RNA sequencing for low nitrogen stress in potato. PLoS ONE 15(5): e0233076. https://doi.org/10.1371/journal.pone.0233076

22. Wan, LC., Zhang, H., Lu, S. et al.,2012. Transcriptome-wide identification and characterization of miRNAs from *Pinus densata*. BMC Genomics 13, 132. https://doi.org/10.1186/1471-2164-13-132

23. Oh T. J., Wartell R. M., Cairney J., Pullman G. S., 2008. Evidence for stage-specific modulation of specific microRNAs (miRNAs) and miRNA processing components in zygotic embryo and female gametophyte of loblolly pine (*Pinus taeda*). New Phytol. 179(1): 67–80. 10.1111/j.1469-8137.2008.02448.x

24. Navarro L., Dunoyer P., Jay F., Arnold B., Dharmasiri N., Estelle M., et al., 2006. A plant miRNA contributes to antibacterial resistance by repressing auxin signaling. Science 312 436–439. 10.1126/science.1126088

25. Gupta OP, Nigam D, Dahuja A, et al., 2017. Regulation of Isoflavone Biosynthesis by miRNAs in Two Contrasting Soybean Genotypes at Different Seed Developmental Stages. Front Plant Sci.; 8 (567). doi:10.3389/fpls.2017.00567

26. Livak KJ, Schmittgen TD., 2001. Analysis of relative gene expression data using real-time quantitative PCR and the 2(-Delta Delta C(T)) Method. Methods.;25(4):402–408. doi:10.1006/meth.2001.1262

27. Hao, C., Xia, Z., Fan, R. et al., 2016. De novo transcriptome sequencing of black pepper (*Piper nigrum* L.) and an analysis of genes involved in phenylpropanoid metabolism in response to *Phytophthora capsic*. BMC Genomics 17, 822. https://doi.org/10.1186/s12864-016-3155-7

28. Xiao-wan Xu, Tao Li, Ying Li, Zhen-xing Li, 2015. Identification and Analysis of *C. annuum* microRNAs by High-throughput Sequencing and their Association with High Temperature and High Air Humidity Stress. Int.J. Bioautomation, 19(4), 459–472

29. Liu Z, Zhang Y, Ou L, Kang L, Liu Y, Lv J, Wei G, Yang B, Yang S, Chen W, Dai X, Li X, Zhou S, Zhang Z, Ma Y, Zou X., 2017. Identification and characterization of novel microRNAs for fruit development and quality in hot pepper (*Capsicum annuum* L.). Gene. Apr 15;608:66–72. doi: 10.1016/j.gene.2017.01.020. Epub 2017 Jan 22. PMID: 28122266.

30. López-Galiano, M.J.; Sentandreu, V.; Martínez-Ramírez, A.C.; Rausell, C.; Real, M.D.; Camañes, G.; Ruiz-Rivero, O.; Crespo-Salvador, O.; García-Robles, 2019. Identification of Stress Associated microRNAs in *Solanum lycopersicum* by High-Throughput Sequencing. Genes, 10(475); doi: 10.3390/genes10070475

31. Cao, X., Wu, Z., Jiang, F., Zhou, R., & Yang, Z., 2014. Identification of chilling stress-responsive tomato microRNAs and their target genes by high-throughput sequencing and degradome analysis. BMC genomics, 15(1), 1130. https://doi.org/10.1186/1471-2164-15-1130

32. Wang Ketao, Su Xiaomei A, Cui Xia A, Du Yongchen A, Zhang Shuaibin A, and Gao Jianchang, 2018. Identification and Characterization of microRNA during *Bemisia tabaci* Infestations in *Solanum lycopersicum* and *Solanum habrochaites*. Horticultural Plant Journal, 4 (2): 62–72.

33. Om Prakash Gupta, Pradeep Sharma, Raj Kumar Gupta, Indu Sharma, 2014. Current status on role of miRNAs during plant-fungus interaction, Physiological and Molecular Plant Pathology; 85, 1–7

34. Zhao JP, Jiang XL, Zhang BY, Su XH., 2012. Involvement of microRNA-mediated gene expression regulation in the pathological development of stem canker disease in Populus trichocarpa. PLoS One; 7(9):e44968.

35. Zhu J, Li W, Yang W, Qi L, Han S. 2013. Identification of microRNAs in *Caragana intermedia* by high-throughput sequencing and expression analysis of 12 microRNAs and their targets under salt stress. Plant Cell Rep.;32(9):1339–1349. doi:10.1007/s00299-013-1446-x

36. Salvador-Guirao, R., Hsing, Y. I., & San Segundo, B. (2018). The Polycistronic miR166k-166h Positively Regulates Rice Immunity via Post-transcriptional Control of *EIN2*. Frontiers in plant science, 9, 337. https://doi.org/10.3389/fpls.2018.00337

37. Chopperla R, Mangrauthia Satendra K, Bhaskar Rao T, Balakrishnan M, Balachandran SM, Prakasam V, Channappa G.,2020. A Comprehensive Analysis of MicroRNAs Expressed in Susceptible and Resistant Rice Cultivars during *Rhizoctonia solani* AG1-IA Infection Causing Sheath Blight Disease. Int J Mol Sci. 27;21(21):7974. doi: 10.3390/ijms21217974. PMID: 33120987; PMCID: PMC7662745.

38. Jin Y, Guo HS., 2018. Plant Small RNAs Responsive to Fungal Pathogen Infection. Methods Mol Biol.; 1848:67–80. doi: 10.1007/978-1-4939-8724-5_6. PMID: 30182229.

39. Li, Y., Zhang, Q., Zhang, J., Wu, L., Qi, Y., & Zhou, J. M., 2010. Identification of microRNAs involved in pathogen-associated molecular pattern-triggered plant innate immunity. Plant physiology, 152(4), 2222–2231. https://doi.org/10.1104/pp.109.151803

40. L. Almagro, L. V. Gómez Ros, S. Belchi-Navarro, R. Bru, A. Ros Barceló, M. A. Pedreño,2009. Class III peroxidases in plant defence reactions, *Journal of Experimental Botany*, Volume 60(2) Pages 377–390.https://doi.org/10.1093/jxb/ern277

41. Passardi F, Penel C, Dunand C., 2004. Performing the paradoxical: how plant peroxidases modify the cell wall. Trends Plant Sci.;9(11):534–540. doi:10.1016/j.tplants.2004.09.002

42. Ricardo B. Ferreira, Sara Monteiro, Regina Freitas, Cláudia N. Santos, Zhenjia Chen, Luís M. Batista, João Duarte, Alexandre Borges and Artur R. Teixeira, 2007. The role of plant defence proteins in fungal pathogenesis. Molecular Plant Pathology; 5, 677–700

43. Hong JK, Jung HW, Kim YJ, Hwang BK., 2000. Pepper gene encoding a basic class II chitinase is inducible by pathogen and ethephon. Plant Sci.;159(1):39–49. doi:10.1016/s0168-9452(00)00312-5

44. Kim, D. S., Kim, N. H., & Hwang, B. K., 2015. The *Capsicum annuum* class IV chitinase ChitIV Interacts with receptor-like cytoplasmic protein kinase PIK1 to accelerate PIK1-triggered cell death and defence responses. Journal of experimental botany, 66(7), 1987–1999. https://doi.org/10.1093/jxb/erv001

45. Kim JH, Woo HR, Kim J, et al., 2009. Trifurcate feed-forward regulation of age-dependent cell death involving miR164 in Arabidopsis. Science.; 323(5917):1053–1057. doi:10.1126/science.1166386

46. Chuck G, and O’Connor D., 2010. Small RNAs going the distance during plant development. Current Opinion in Plant Biology, 13: 40–45.

47. Zhang, H. X., Feng, X. H., Ali, M., Jin, J. H., Wei, A. M., Khattak, A. M., & Gong, Z. H. (2020). Identification of Pepper *CaSBP08* Gene in Defense Response Against *Phytophthora capsici* Infection. Frontiers in plant science, 11, 183. https://doi.org/10.3389/fpls.2020.00183

48. Zhang HX, Ali M, Feng XH, Jin JH, Huang LJ, Khan A, Lv JG, Gao SY, Luo DX, Gong ZH., 2018a. A Novel Transcription Factor *CaSBP12* Gene Negatively Regulates the Defense Response against *Phytophthora capsici* in Pepper (*Capsicum annuum* L.). Int J Mol Sci. 22;20(1):48. doi: 10.3390/ijms20010048. PMID: 30583543; PMCID: PMC6337521.

49. Zhang HX, Feng XH, Jin JH, Khan A, Guo WL, Du XH, Gong ZH., 2020b. *CaSBP11* Participates in the Defense Response of Pepper to *Phytophthora capsici* through Regulating the Expression of Defense-Related Genes. Int J Mol Sci. 28;21(23):9065. doi: 10.3390/ijms21239065. PMID: 33260627; PMCID: PMC7729508.

